# Hox genes modulate physical forces to differentially shape small and large intestinal epithelia

**DOI:** 10.1101/2023.03.15.532602

**Authors:** Hasreet K. Gill, Sifan Yin, Nandan L. Nerurkar, John C. Lawlor, Tyler R. Huycke, L. Mahadevan, Clifford J. Tabin

## Abstract

The small and large intestines arise from a common primordial gut tube but ultimately become specialized in both form and function. While the midgut forms villi, the hindgut develops flat, brain-like sulci that resolve into heterogeneous outgrowths. Gut compartment identities are demarcated early in development via Hox genes, which are highly conserved, master regulators of spatial patterning in the embryo. Yet, how these factors trigger regional morphogenesis has remained a mystery. Combining mechanical measurements and mathematical modeling, we demonstrate that the posterior Hox gene *Hoxd13* regulates biophysical phenomena that shape the hindgut lumen. We further show that *Hoxd13* acts through the TGFβ pathway to thicken, stiffen, and promote isotropic growth of the subepithelial mesenchyme; together, these features generate hindgut surface patterns. TGFβ, in turn, promotes collagen deposition to affect mesenchymal geometry and growth. We thus identify a cascade of events downstream of genetic identity that direct posterior intestinal morphogenesis.

## INTRODUCTION

The Hox genes are transcription factors that play critical roles in organizing developmental patterning across metazoa, particularly along the anterior-posterior body axis. They were first identified in Drosophila, where mutations in these genes lead to homeotic transformations of body segments (Lewis 1963). Further genetic analyses led Ed Lewis to propose that, in the fly, Hox genes form a combinatorial code, specifying the identity and morphogenesis of the arthropod segments (Lewis, 1978). The subsequent finding of a highly conserved domain within these genes, the homeobox (McGinnis, Levine, et al., 1984; Scott & Weiner, 1984), led to the realization that the Hox gene family arose and expanded through tandem duplication. Just a few months after this discovery, it was found that highly related genes also exist in vertebrates and other metazoa (Carrasco et al., 1984; McGinnis, Garber, et al., 1984).

Just as in Drosophila, tetrapod Hox genes play a central role in controlling the development of the body plan. 39 genes are arranged into 4 clusters, and within each cluster, the chromosomal order of the Hox genes matches the order of their expression domains in the developing embryo. Based on knowledge of Hox gene expression patterns in the mouse, in 1993, Michael Kessel and Peter Gruss proposed that the overlapping pattern of Hox gene activity might serve as a code for establishing differential vertebral morphologies, in analogy to the roles their Drosophila counterparts play in determining segment identity (Kessel & Gruss, 1991). Since that time, extensive loss of function analyses in mice have verified that this model is essentially correct (reviewed in Wellik, 2007). It is striking, however, that more than 30 years since Kessel and Gruss put forth the idea of a Hox vertebral code, we still have little idea of how it works. For example, an anatomist can immediately tell the difference between cervical (neck) and lumbar (abdominal) vertebrae, and the developing cervical vertebrae express Hox 2, 3, 4, and 5 paralogs, while lumbar vertebrae express Hox 7, 8, and 9 paralogs. But how these different groups of Hox genes direct the differences in vertebral morphology—namely, the downstream genetic and/or cellular process they act through—remains a black box. In fact, though studies in Drosophila have uncovered Hox-regulated cellular and mechanical phenomena that drive organogenesis, such mechanistic knowledge is virtually nonexistent for vertebrate organ systems (De Las Heras et al., 2018; Mitchell et al., 2022).

While axial skeletal morphology is the classic example of anterior-posterior Hox patterning in vertebrates, Hox genes contribute to the differential anterior-posterior patterning of a variety of embryonic tissues, from the central nervous system (reviewed in Parker & Krumlauf, 2020), to the reproductive tract (reviewed in Major et al., 2022). An additional organ system where Hox genes have been implicated in this context is the developing gastrointestinal (GI) tract. The GI tract initially forms as a morphologically uniform primitive gut tube, encompassing the foregut, midgut and hindgut. Over the course of embryogenesis, this tube differentiates into the distinct organ structures required for digestion and the absorption of nutrients, such as the pharynx, esophagus, stomach, small intestine, and large intestine. Each of these GI compartments has a unique function, and correspondingly displays distinct morphological features. For example, the lumen of the small intestine is characterized by long finger-like epithelial projections called villi, while the lining of the large intestine has much squatter and superficial epithelial folds that have been referred to as “colonic villi”, or epithelial cuffs (Bell & Williams, 1982; De Santa Barbara et al., 2003; Mahoney et al., 2008).

Shortly after the discovery of the vertebrate Hox clusters, it was observed that the Abdominal-B related Hox genes (Hox paralog groups 9 – 13) are expressed in a nested set of overlapping domains with distinct regional boundaries demarcating domains of the hindgut and midgut (Roberts et al., 1995; Yokouchi et al., 1995). For example, Hox 13 paralogs are expressed in the terminal end of the hindgut in humans, mice, chicks, zebrafish, flies, worms, and sea urchins (Annunziata et al., 2019; Kondo et al., 1996). *Hoxd13* loss-of-function destroys the muscular apparatus of the anal sphincter in the mouse, and infection of the chick gut with RCAS retrovirus carrying a dominant negative form of *Hoxa13* causes hindgut and cloacal atresia (Barbara & Roberts, 2002; Dan et al., 2010; Warot et al., 1997; Zákány & Duboule, 1999). Most striking, however, is the finding that *Hoxd13* overexpression in the chick midgut is sufficient to replace villi with an epithelial lining that resembles that of the hindgut and produce the acid mucin-secreting cells typical of the large intestine (Roberts et al., 1998). Hoxd13 is, thus, both necessary and sufficient to determine critical aspects of hindgut specific morphology—but, how it accomplishes this has, once again, remained a black box.

While the mechanisms underlying differences in morphogenesis of the distinct functional domains of the gut have remained unresolved, significant progress has been made in understanding the morphogenesis of the epithelium of the midgut (Shyer et al., 2013). The villi of the chick small intestine forms through a stepwise process, in which the gut epithelium and subjacent mesenchyme first fold into longitudinal ridges, which then buckle into a zigzag pattern, before finally distorting into ordered arrays of villi. This process is orchestrated by the sequential differentiation of distinct smooth muscle layers, the first oriented circumferentially and the latter two orientated longitudinally.

These act to restrict the expansion of the growing epithelium and mesenchyme, creating the compressive stresses that lead to their buckling. For example, when the first circumferential smooth muscle layer differentiates, it prevents radial expansion of the growing inner layers of the gut. With continued proliferation, these layers are forced to buckle inward, as they cannot expand outward. The additional differentiation of the second longitudinal layer of smooth muscle leads to biaxial compression and zigzags, followed by a third longitudinal layer that, along with additional cellular mechanisms, further compresses zigzags into primordial villi.

As informative as these findings are in understanding the stepwise morphogenesis of villi in the chick small intestine, the model does raise a conundrum, as the same three smooth muscle layers arise in the same sequence in the hindgut (Gill, Yin et al. 2023). Hence, the hindgut presumably faces very similar biophysical constraints, yet it achieves a very different epithelial morphology. One possible answer to this would lie in different physical characteristics of the two developing gut segments, as the way a tissue deforms when constrained depends on its mechanical properties and differential growth-induced strains. This begs the question of how, in fact, the properties of the developing midgut and hindgut differ physically, and whether the differences can indeed explain differential epithelial morphogenesis. Here, we identify how distinct mechanical and growth parameters in the midgut and hindgut give rise to their unique morphologies. We further uncover that these regional tissue mechanics are specified by upstream Hox genes, which pattern the tissue through TGFβ signaling.

## RESULTS

### Distinct stepwise morphogenesis of the epithelia in the midgut and hindgut

Villi in the midgut display crystalline regularity in their shapes and arrangement (Coulombre & Coulombre, 1958). As we and others have previously described (Shyer et al., 2013), villi arise in a stepwise manner wherein the smooth luminal lining of the midgut at embryonic day (E) 8 transforms into a series of ridges at E12, zigzags at E14 and protovilli—bulges that prefigure final villi locations, born from in-plane rotations of zigzag arms—by E18 (Figure 1A-B). In contrast, the first observable deformation of the luminal surface of the hindgut occurs at E12 when it forms superficial, brain-like sulci, which become extensively branched by E14 and evolve into cuffs by E16 (Figure 1A). Though tightly packed like primordial villi, cuffs are shorter, wider, heterogeneous in size, and appear spatially disordered (Figure 1A, E).

**Figure 1.**
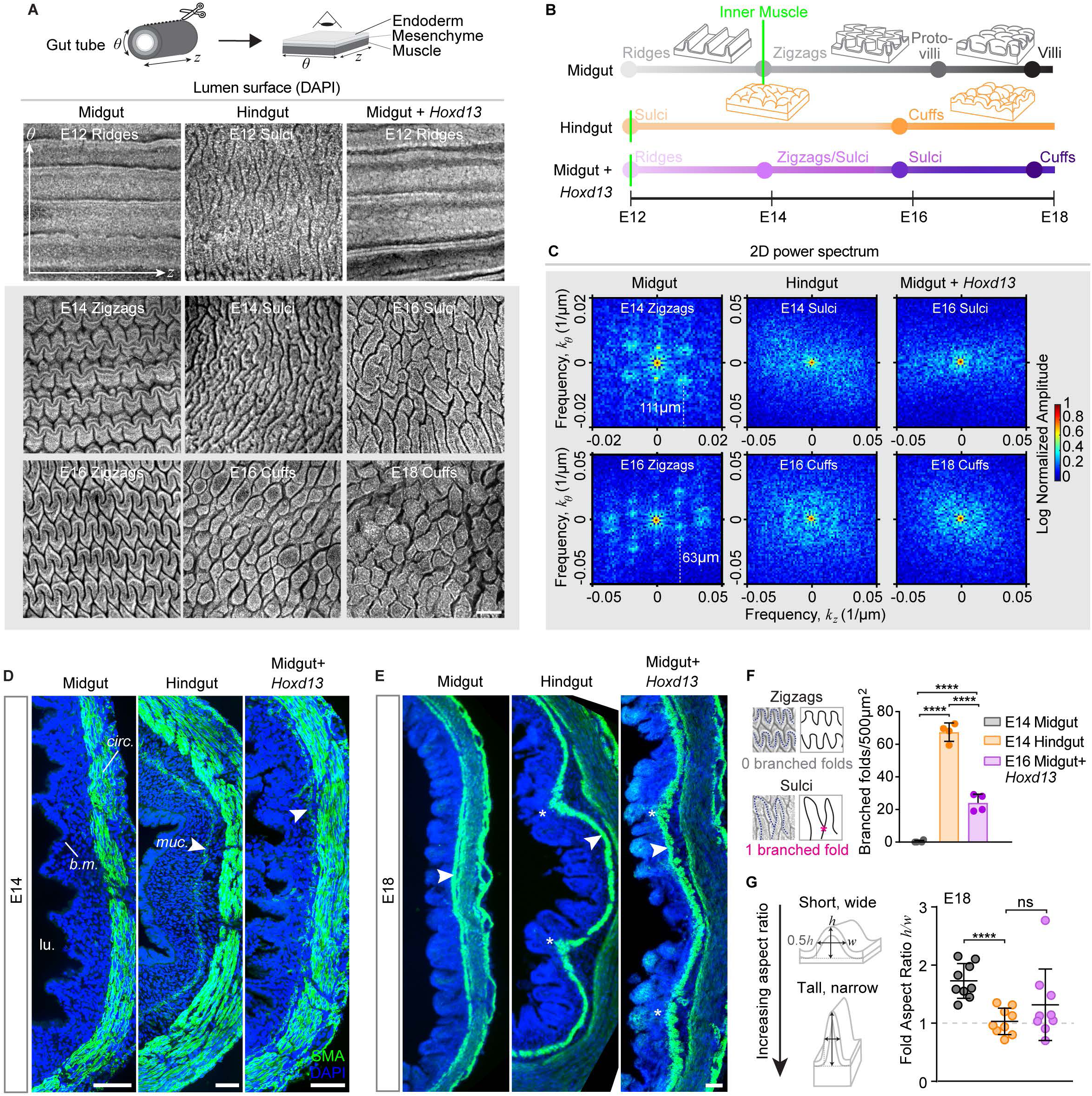
The chick hindgut adopts distinct lumen wrinkling patterns from the midgut, which are replicated with Hoxd13 misex­ pression. (A) Notation for gut axes and method of imaging the surface of the endoderm after cutting the tube open longitudinally. Lumen surface morphologies of midgut, hindgut, and RCAS-Hoxd13 midgut (“Midgut + Hoxd13”) at E12 for all conditions, E14 and E16 for the midgut and hindgut, and E16 and E18 for the midgut + Hoxd13. Scale bar, 100µm. (B) Timeline of lumen morphogenesis in the midgut, hindgut and midgut + Hoxd13, with cartoon representation of key morphologies. The shallowness/steepness of each color gradient represents the timescale of the morphological transition. Green lines indicate muscularis mucosa differentiation. (C) Power spectral density profiles corresponding to spatial domain images within the gray box in (A). Color indicates log normalized amplitude. For patterns with peaks corresponding to characteristic pattern wavelengths, wavelength values on the longitudinal axis (z) are indicated on the plot. (F, E) Trans­ verse sections at E14 (E) and E18 (F) depicting one hemisphere of the control midgut, hindgut, and RCAS-Hoxd13 midgut. SMA, Alpha Smooth Muscle Actin; Calp, Calponin 1; b.m. and straight line, basement membrane; lu., lumen; circ., circumferential muscle layer; muc. and arrowhead, muscularis mucosa; asterisk, secondary large-scale fold. Scale bar, 50µm. (F) Method of identifying branch points in lumen patterns by tracing the peaks of folds from raw lumen surface images. Number of branched folds per 500µm2 at E14 for the midgut and hindgut, and E16 for the RCAS-Hoxd13 (****, p<0.0001; t-test, n=4 images). (G) Method of measuring fold aspect ratios as height divided by width, where width is measured at the fold half max; tall, thin folds have larger values and short, wide folds have smaller values. Aspect ratios of single folds at E18 only(****, p<0.0001; ns, not significant, t-test, n=10 folds).

Concomitant with the maturation of sulci in the E14 hindgut, a second order buckling that includes both the endoderm and the inner layer of mesenchyme appears (Figure 1D; Gill, Yin et al. 2023). As this is also the time when the innermost layer of longitudinal smooth muscle (muscularis mucosa) differentiates, which forms in a pattern of periodic deformations distanced from the subjacent circumferential smooth muscle— a configuration maintained as the sulci give way to cuffs (Figure 1E). By comparison, in the midgut, there is no such second-order folding and the circumferential layer of smooth muscle forms as a flat band beneath the epithelial folds, sandwiched between the closely apposed outer and inner longitudinal smooth muscle layers (Figure 1E).

It is striking that the same three smooth muscle layers emerge sequentially in the hindgut, in the same order, and with the same fiber orientation as the midgut (Gill, Yin et al. 2023). Indeed, given that their role in midgut epithelial morphogenesis is to provide resistance to directional expansion, leading to stresses that buckle the inner layers into ridges and then zigzags, the fact that muscles are thicker in the hindgut and that each differentiates 1-2 days earlier should only make them more effective in driving such morphological transformations (Figure 1B, D). The fact that they do not do so suggests that differences in the geometric and mechanical properties of the inner layers themselves must differ between the midgut and hindgut, thus affecting how regional epithelia interpret physical constraints from muscle.

### The developing hindgut and midgut have distinct physical properties

Clues into the different modes of buckling in the midgut and hindgut come from extensive previous theoretical analyses of the formation of sulci, which superficially resemble the reticulated grooves that first appear on the luminal surface of the hindgut (Hohlfeld & Mahadevan, 2011, 2012; Tallinen et al., 2013). Sulci are unique, nonlinear surface instabilities that have recently been studied in the context of brain, though they appear widely in nature. Previous work would suggest that sulci in the gut may emerge when the endoderm-mesenchyme bilayer is compressed biaxially (through transversely isotropic expansion), where the modulus ratio of the two layers is low (i.e., they are equally stiff, or the endoderm layer is softer) (Ciarletta et al., 2014; Holland et al., 2018; Razavi et al., 2016; Tallinen et al., 2014; Tallinen & Biggins, 2015). To test this possibility, we measured differential growth, geometric features, and stiffnesses of inner endoderm and mesenchyme layers during late-stage midgut and hindgut lumen morphogenesis.

To measure strain in these layers resulting from their growth, we dissected apart the composite of inner epithelium and mesenchyme (down to the level of the first smooth muscle layer). We then allowed layers to reach their stress-free lengths and determined strain as the percent increase or decrease in tissue length in the relaxed configuration (Figure 2A) (Fung & Liu, 1989; Shyer et al., 2013; Zhao et al., 2007; Gill, Yin et al., 2023). Measurements from both circumferential and longitudinal dissections yielded differential growth profiles over time. As expected, the midgut initially accrues strain only circumferentially (negative values in Figure 2B), reflecting the circumferential compression that forces the flat lumen to form ridges. Subsequently, between E12 and E14, longitudinal compression appears, concomitant with differentiation of the outer muscle layer, while at this stage the circumferential strain no longer increases (Figure 2B). Therefore, within each growth period (E8-E12, E12-E14, E14-E18), midgut growth is highly anisotropic, as previously reported (Amar and Jia 2013). By stark contrast, but consistent with the theoretical analyses of sulci formation, the hindgut gradually develops compression on both axes nearly simultaneously, resulting in equibiaxial growth during most of lumen morphogenesis (Figure 2B).

**Figure 2.**
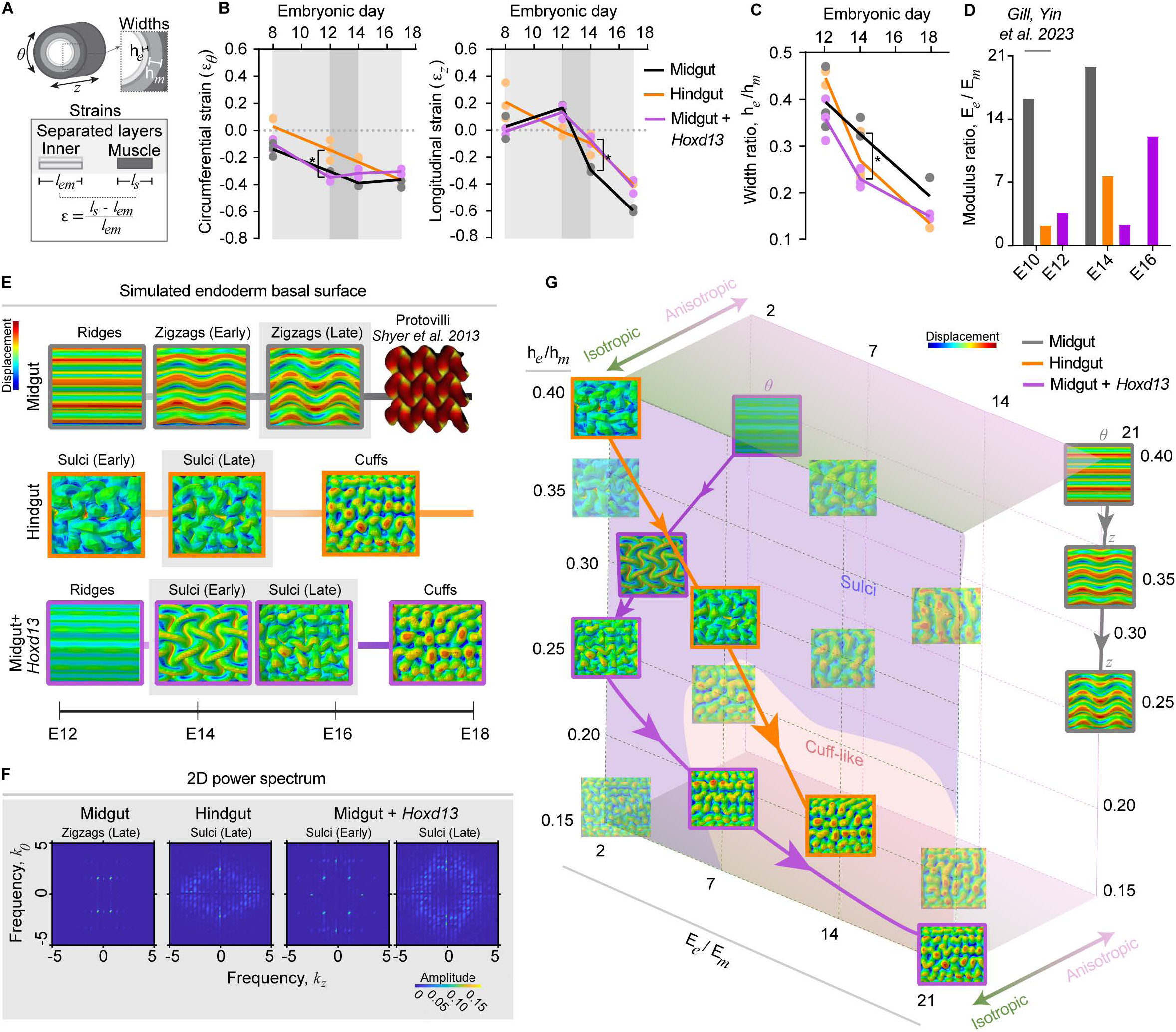
Hindgut geometric, material, and growth properties diverge from those of the midgut and are promoted by Hoxd13 activity. (A) Measurement of widths and strains from cross-sections and dissected, relaxed inner and outer muscle layers (s), respectively. Strain is determined as the percent change in length of inner layers upon constraint by muscle. (B) Differential growth properties of the midgut, hindgut, and RCAS-Hoxd13 midgut (“Midgut + Hoxd13”). Gray boxes in each plot frame time periods of interest, where dark gray corresponds to the biaxial buckling transition. Left, Circumferential (8) and right, longitudinal (z) strains, *e,* over time. Dotted gray lines at 0.0 strain in each plot indicate transition between inner composite tensile (>0) and compressive (<0) regimes(*, p<0.05, t-test, n=2-3). (C, D) Ratios of endoderm (e) to mesenchyme (m) (C) width, h (*, p<0.05, n=3), and (D) Young’s modulus, E, over time. Midgut and hindgut modulus ratios at E10 are used here for comparison to the RCAS-Hoxd13 midgut at E14. (E) Numerical simulation results at successive stages of lumen morphogenesis shown in Figure 1B. In a tube geometry, views correspond to the outer surface of the endoderm layer for ease of visualizing the morphology. Color indicates magnitude of displacement. (F) 2D power spectral densities for major wrinkling modes indicated by gray boxes around simulation results in (D). (G) 3-dimensional parameter space representing experimental lumen folding trajectories (lines and arrows, which indicate progression over time) and simulation results (boxes) used to identify two morphological domains (blue “sulci” and orange “cuff-like” colored regions) as functions of modulus ratio, growth, and thickness ratio. For cases with anisotropic growth, 8 and z above the boxes represent dominating growth directions.

Next, tissue width measurements taken from transverse sections (Figure 2A) revealed that both the endoderm and mesenchyme layers are thicker in the hindgut (Figure S3B, D). Yet, the ratio of endoderm to mesenchyme width is not different until the appearance of sulci from E12-E14, when the width ratio sharply drops in the hindgut as the mesenchyme thickens (Figure 2C, S3C).

Given that E12-E14 is the key time frame when the midgut and hindgut develop distinct biaxial morphologies, we next measured Young’s moduli of the endoderm and mesenchyme layers at E14. To do so, we used a custom uniaxial tensile testing system developed to compare regional physical properties several days prior, during buckling initiation along the foregut, midgut and hindgut (described in detail in Gill, Yin et al., 2023; Nerurkar et al., 2017) (Figure S3A). We first observed that, consistent with measurements during uniaxial buckling at E10, the endoderm is softer in the hindgut at E14 (Figure S3B; Gill, Yin et al., 2023). By contrast, mesenchyme modulus, calculated from average endoderm and composite modulus and width measurements, is higher in the hindgut at E14; together with a softer endoderm, this lowers the modulus ratio, again consistent with E10 data (Figure 2D, S3B-C). Thus, at the critical time when the midgut forms zigzags, but the hindgut forms sulci, the modulus ratio between the endoderm and mesenchyme is high in the midgut and low in the hindgut (Figure 2D). Notably, the modulus ratio in the hindgut trends upward as the sulci gives way to cuffs (Figure S3D).

Taken together, the elastic, geometric, and growth properties of the hindgut are both distinct from those of the midgut and qualitatively consistent with theoretical work on the emergence of creased sulci patterns.

### A mathematical model incorporating measured physical parameters recapitulates midgut and hindgut morphogenesis

Despite similarities between our observations and theoretical expectations, determining whether measured values sufficiently explain different lumen patterns required turning to computational modeling. We therefore constructed a mathematical model based on a widely used differential growth framework (Shyer et al., 2013; Ciarletta et al., 2014; Tallinen & Biggins, 2015), but now combined with experimentally measured spatiotemporal physical parameters.

To simulate late-stage ridge, zigzag, sulci, and cuff formation, we first adapted our two-dimensional growing bilayer model of early buckling initiation to three dimensions (Methods; Gill, Yin et al., 2023). We therefore applied volumetric growth theory as before, with the deformation gradient decomposed as 𝑭 = 𝑨 ⋅ 𝑮. 𝑨 is the elastic deformation tensor and 𝑮 = diag(𝑔_𝑟_, 𝑔_𝜃_, 𝑔_𝑙_) is a spatiotemporal growth tensor capturing the addition of material volume and the plastic (irreversible) deformation in three dimensions. Treating the gut as a Neo-Hookean elastic solid defines strain density as

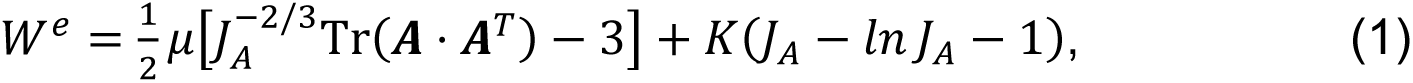

where 𝜇 and 𝐾 are the initial shear modulus and bulk modulus, respectively; *J_A_* = *det*(***A***) is the elastic volumetric strain. Numerical simulations were performed using the finite element method (Methods).

Fitting the spatiotemporal growth tensor to the differential growth behaviors of the midgut and hindgut—stepwise anisotropic growth and transversely isotropic growth, respectively—yielded two representative sets of functions (Methods, Table S1).

Combined with measured thickness and modulus ratios at each stage of lumen wrinkling, these growth tensors predicted the distinct morphological trajectories of the midgut and hindgut. Monotonically increasing and decreasing longitudinal and circumferential growth profiles, respectively, in the context of a relatively high modulus ratio and moderately decreasing thickness ratio yields ridges followed by zigzags (Figure 2E). By contrast, a low modulus ratio with linearly increasing longitudinal and circumferential growth results in sulci at both high and low thickness ratios (Figure 2E, G). Further exploration of the parameter space defined by these three characteristics revealed that, indeed, sulci appearance is driven by transversely isotropic growth, though properties of the folds vary with modulus and thickness ratios—for example, sulci wavelength appears to scale with thickness ratio (Figure 2G).

Most striking from our parameter space, however, was the observation that in the high modulus ratio and low thickness ratio regime, the surface pattern becomes limited to spatially segmented outgrowths resembling cuffs instead of labyrinthine sulci (Figure 2G). Given the trend of increasing modulus ratio and drastically decreasing width ratio in the hindgut, its morphological trajectory ultimately lands in this predicted “cuff-like” state (Figure 2C-E, G, S3D). Thus, unlike in the midgut where the final stages of villus outgrowth cannot be explained by a continuum-level mechanical model of lumen wrinkling (Shyer et al., 2013), our model captures all major steps of hindgut morphogenesis from biophysical parameters alone, and further indicate that these properties are sufficient to drive luminal morphogenesis in the hindgut.

### *Hoxd13* misexpression in the midgut transforms luminal folding a hindgut-like pattern

With a biomechanical understanding of chick large and small intestinal lumen wrinkling in hand, we next turned our focus to the genetic regulation of these patterns. At the time of hindgut morphogenesis, *Hoxd13* is expressed in the developing hindgut in a domain extending more proximally through the hindgut than previously reported at earlier stages (Figure S1). When misexpressed in the developing midgut, *Hoxd13* transforms the luminal folding pattern into a phenocopy of the normal hindgut (Roberts et al., 1998). To assess this phenotype in more detail, a replication competent retroviral vector transducing *Hoxd13* (RCAS-*Hoxd13*) was electroporated into the midgut lateral plate mesoderm at E2.5 (Methods; Roberts et al., 1995). At early stages of gut morphogenesis, the infected midguts look indistinguishable from controls (Figure 1A).

However, when the hindgut forms sulci and the midgut adopts a zigzag configuration at E14, the *Hoxd13*-expressing midgut displays an intermediate phenotype between the two (Figure 1A, S2A). By day 16, the *Hoxd13*-expressing midgut ridges have transformed to a sulci pattern like that of the day 14 hindgut, albeit with a two-day delay (Figure 1A). By day 18, instead of an array of thin and long primordial villi, the *Hoxd13*-expressing midguts have formed short, and wide cuffs like those normally seen in the day 16 or 18 hindgut (Figure 1A, G)

Moreover, the E18 *Hoxd13*-expressing midguts display the secondary buckling characteristic of the hindgut at this stage, wherein the innermost layer of smooth muscle forms periodic folds along with the adjacent mesenchyme, as in the hindgut (Figure 1E). Radial quantifications of smooth muscle actin (SMA) stain intensity from the basement membrane to circumferential muscle illustrate the shift in the positions of smooth muscle peaks away from the muscle boundary in these conditions (Figure S2D). Also, as noted above, the smooth muscle layers differentiate earlier in the wild-type hindgut than midgut. In the *Hoxd13*-expressing midgut, the innermost longitudinal layer has already formed at E14, recapitulating earlier muscle differentiation of the hindgut (Figure 1B, D).

Taken together, at least superficially, Hoxd13 activity appears to be sufficient to posteriorize midgut folding and smooth muscle differentiation. To quantitatively determine if this is the case, we assessed properties of wrinkling patterns in the midgut, hindgut, and *Hoxd13*-expressing midgut; comparing the E16 RCAS-*Hoxd13* midgut (when it appears to form sulci) to E14 midgut (zigzags) and hindgut (sulci); and the E18 RCAS-*Hoxd13* midgut (when it appears to form cuffs) to the E16 midgut (late zigzags) and hindgut (cuffs) (Figure 1A-B).

First, we used a metric assessing the extent of branching of the epithelial folds (Methods, Figure 1F). While zigzags in the E14 midgut are essentially unbranched, we observe significant branching in both the E14 hindgut and E16 *Hoxd13*-expressing midgut (Figure 1F). At later stages, we used Delaunay triangulation to assess the relative presence of cuff vs. pre-villi outgrowths. In line with our observations, both hindgut cuffs and *Hoxd13*-expressing midgut cuffs show higher variance in their Delaunay triangle edge length distributions compared to primordial villi, consistent with relatively higher spatial disorder in the surface arrangements of outgrowths (Figure S2C). A parallel analysis of Voronoi cell area distributions supported this result.

To further assess the pattern and degree of order in the epithelial folds, we examined the 2D Fast Fourier Transformation (FFT) power spectra, again comparing E16 *Hoxd13*-expressing midguts to wild type E14 midguts and hindguts, and E18 *Hoxd13*-expressing midguts and wild type E16 midguts and hindguts. In the normal midgut, the 2D FFT plot at each time point shows a discrete set of peaks at the characteristic pattern wavelengths, where the longitudinal period length decreases as the herringbone pattern is compressed with time (Figure 1C). Complementary autocorrelation profiles separating patterns on the circumferential (*ϴ*) and longitudinal (*z*) axes reveal high-amplitude sinusoidal waves across time points (Figure S2B). Together, these measures capture the stereotyped periodicities of ridge, zigzag and primordial villi patterns.

In contrast, the hindgut has a diffuse spectral density profile at all stages and lower autocorrelation amplitudes, though slight longitudinal bias appears at later stages (Figure 1C, S1B). Accordingly, autocorrelation plots and amplitudes present only moderate evidence of periodic wrinkling on the longitudinal axis, as well as the large-scale circumferential buckling not seen in the midgut (Figure S2B). The E16 and E18 *Hoxd13*-expressing midguts, strikingly, display 2D power spectra and autocorrelation profiles like those of the hindgut two days earlier in each case (Figure 1C, S2B). This analysis therefore supports the contention that ectopic expression of *Hoxd13* drives the midgut towards a hindgut-like morphology—namely, branched wrinkling and diminished pattern periodicity.

### Hoxd13 modulates mechanical properties of the developing gut tube

To test whether Hoxd13 alters luminal gut morphogenesis by modulating the physical parameters we identified as distinguishing the midgut from the hindgut, we again measured differential growth, stiffness, and geometric features of inner endoderm and mesenchyme layers, but now in *Hoxd13*-expressing midguts. Consistent with it initially forming midgut-like ridges, growth of the *Hoxd13*-expressing midgut is indistinguishable from that of the normal midgut until E12 (Figure 2B). However, the dramatic increase in longitudinal compression that normally forces buckling into zigzags between E12 and E14 in the midgut is dampened with Hox misexpression, making differential growth less anisotropic during this period. Subsequent differential growth trajectories are closer to that of the hindgut, suggesting, once again, that the biaxial buckling transition is the key time point when the normal and transformed midgut patterns diverge, and that this deviation in the *Hoxd13*-expressing midgut pattern evolution is marked by loss of anisotropic growth.

We next addressed thickness and modulus ratios between growing layers in the composite endoderm and mesenchyme and found that, like the hindgut, the *Hoxd13*-expressing midgut mesenchyme is notably thicker than that of the normal midgut at the time that their morphological trajectories diverge, which lowers the width ratio (Figure 2C, S3C). The composite modulus of the *Hoxd13*-expressing midgut is significantly higher, corresponding to a stiffer mesenchyme, and consequently a lower modulus ratio relative to the endoderm—in fact, the ratio at E14 is identical to the hindgut modulus ratio just a few days prior (Figure 2D, S2B-C). Therefore, we conclude that both thickening and stiffening of the mesenchyme, without changes to endoderm properties, is sufficient to imbue the RCAS-*Hoxd13* midgut with hindgut-like physical and geometric characteristics.

To verify that the physical parameters altered by *Hoxd13* misexpression can explain the hindgut-like phenotype we observed, we examined the pattern trajectory predicted by our computational model. We first observed that a transition to isotropic growth with a low modulus ratio, analogous to the transitional E14 morphology in Figure S2A, generated an ordered pattern of sulci from subtle initial ridges (Figure 2E, G).

Subsequent models generated sulci like those of hindgut simulations. As in the hindgut, the *Hoxd13-*expressing modulus ratio trends upward afterward, which, in conjunction with a lower width ratio at E18, leads to segmentation of sulci into cuffs (Figure 2D, G, S3D). Key pattern predictions pulled from the path of each condition across parameter space were quantified using FFT: zigzag patterns show peaks at characteristic wavelengths, while sulci and cuff power spectra are diffuse and circular, all consistent with experimental data (Figure 2F, 1D).

We conclude, therefore, that the breakdown of epithelial ridges into sulci by Hoxd13 is largely achieved through thickening and stiffening of the developing gut mesenchyme—rendering lower modulus and thickness ratios between the endoderm and mesenchyme—and a shift to isotropic growth; cuffs then resolve through a subsequent increase in modulus ratio and decrease in thickness ratio. However, though this finding advances our understanding of the role of Hoxd13 in the hindgut, the mechanism by which this Hox transcription factor alters tissue material and geometric properties remained unclear.

### TGFβ signaling is upregulated in hindgut and *Hoxd13*-expressing midgut subepithelial mesenchymal cells

To begin investigating pathways that might be regulated by Hoxd13 in the process of establishing gut morphology, we performed bulk RNA sequencing to identify genes commonly differentially expressed in sulci/cuff-forming lumens of both the developing hindgut and *Hoxd13*-expressing midgut. Our pattern analysis and mechanical data suggested that E12 to E14 is the critical time when folding instabilities become distinct, so we collected tissue from the midgut, hindgut, and *Hoxd13*-expressing midgut at these times. Also, because *Hoxd13* misexpression was restricted to the mesoderm and mechanical changes occurred mainly in the mesenchyme layer, we only harvested mesodermal RNA (Figure S3B-C, Methods).

Filtering for genes differentially expressed in both the hindgut and *Hoxd13*-expressing midgut relative to the normal midgut yielded a list of 128 genes at E12 and 721 genes at E14, with 62 in common between time points (Figure S4A). Gene function analysis revealed that 16/62 (25%) of these genes are involved at various levels in the TGFβ superfamily—including diffusible ligands and antagonists (*Inhba, Grem1, Chrdl2*), extracellular matrix (ECM) degradation or assembly factors (*Mmp2, Thbs2, Fmod*), downstream targets (*Ptn, Cd44, Actn1*), and others (Figure 3A-C, S4B-C) (Godwin et al. 2019; Sengle and Sakai 2015; Yin et al. 2019). Additional key pathway ligands (*Gdf3, Tgfb1*) and ECM factors (*Mfap5, Mfap2, Col1a2, Col1a1, Eln*), become newly up-or downregulated at E14, when a phenotypic difference is first apparent in the *Hoxd13*-misexpressing midgut (Figure 3B, C) (Penner et al. 2002). The similarity in genes up- and down-regulated in the wildtype hindgut and *Hoxd13*-expressing midgut provides further verification that this Hox misexpression has the effect of posteriorizing the midgut (Figure 3C, S3A).

**Figure 3.**
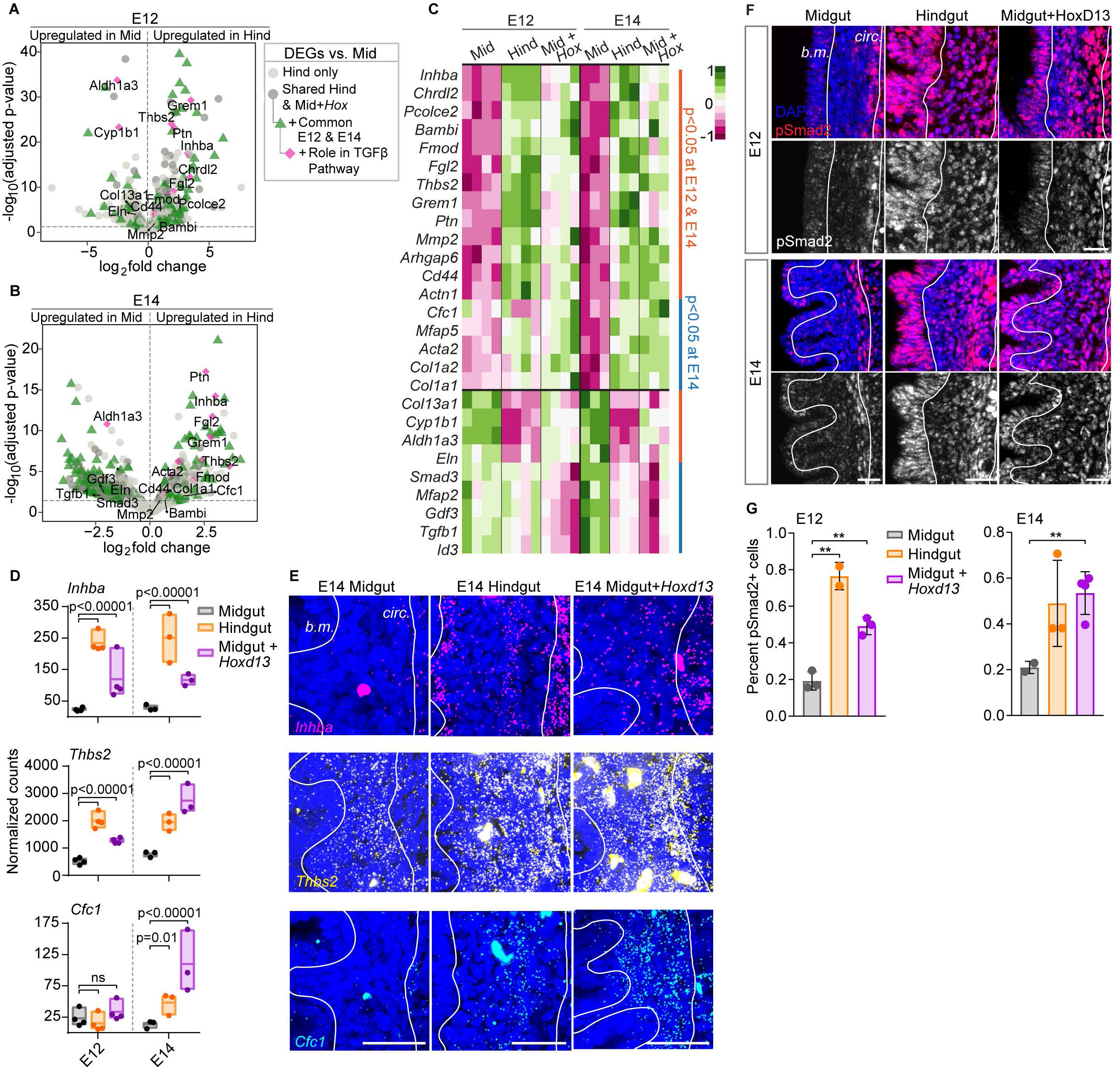
TGFf3 signaling is upregulated in the hindgut and RCAS-Hoxd13 subepithelial mesenchyme. (A, B) Volcano plots with genes differentially expressed between the hindgut and midgut mesodermal layers at (A) E12 and (B) E14. Dashed gray lines indicate the significance cutoff at adjusted p=0.05. Of genes also up- or down-regulated in the RCAS-Hoxd13 midgut vs. control midgut (darker gray dots), a proportion were commonly differentially expressed at both points (green triangles), and a subset of those were genes known to be involved in the TGF pathway (pink diamonds and gene name labels). (C) Heat map of final subset of TGF pathway genes, with genes upregulated in the hindgut and midgut + Hoxd13 in the upper portion, and downregulated below. Significantly differentially expressed genes at both E12 and E14 are indicated by orange lines, and only at E14 by blue lines. (D) Normalized counts with indicated p adjusted values at E12 and E14 for three TGF genes of interest (ns, not significant; n=4 at 12, n=3 at E14) (E) HCR-FISH labeling each gene in the E14 subepithelial mesenchyme bounded by white lines. Note that large, irregular bright spots are auto-fluorescent enteric neural crest cells. Scale bar, 50µm. (F) pSmad2 immunostains in the subepithelial mesenchyme at E12 and E14. b.m., basement membrane; circ., circumferential muscle layer; Scale bar, 20µm. (G) Proportion of pSmad2-positive cells in regions bounded by white lines in (F) for E12 and E14 (**, p<0.01; t-test, n=2-3).

To characterize the expression patterns of the most relevant and significantly upregulated TGFβ pathway genes in our dataset, we performed HCR (hybridization chain reaction) smFISH at E14, focusing on factors that span different tiers of pathway regulation: 1) *Inhba*, a subunit for the TGFβ ligand activin, 2) *Thbs2*, a thrombospondin that is known to release latent TGFβ ligand sequestered in the ECM to promote signaling, and 3) *Cfc1*, a Nodal co-receptor (Figure 3D-E, S4C) (Derynck and Budi 2019; Gurdziel et al. 2016; Havis et al. 2016; Tzavlaki and Moustakas 2020; Wang et al. 2016). *Inhba* and *Thbs2* were differentially upregulated at both time points, while *Cfc1* was upregulated at E14 only (Figure 3D). HCR-FISH revealed that all three genes show increased expression specifically in the subepithelial mesenchyme (Figure 3E). Plots of radial intensity, from the basement membrane to the circumferential muscle, support higher overall expression levels extending to the border mesenchyme-epithelium border (Figure S4D). Therefore, key TGFβ factors are differentially expressed in the same sub-compartment (mesenchyme) where hindgut and *Hoxd13*-expressing midgut physical properties differ from the normal midgut.

To determine whether enrichment for TGFβ genes in our RNA-seq dataset corresponded to differences in pathway activation, we stained for phosphorylated Smad2 (Figure 3F) (Derynck and Budi 2019; Tzavlaki and Moustakas 2020). At both E12 and E14, we observed higher proportions of pSmad2-positive nuclei in mesenchymal cells of the hindgut and *Hoxd13*-expressing midgut and, therefore, concordance between the site of pathway activation and gene upregulation (Figure 3G). Also consistent with HCR results, quantifications of radial pSmad2 signal intensity in this region reveal that, though all three conditions show highest intensities in the innermost longitudinal muscle layers, only segments with posterior identity have pSmad2+ nuclei extending throughout the subepithelial mesenchyme (Figure S4E). As a result, TGFβ activity is not only higher, but also decreases to a lesser extent from the circumferential muscle to basement membrane in the hindgut and *Hoxd13*-expressing midgut.

Together, these results demonstrate that *Hoxd13* expression in the gut is sufficient to induce mesenchymal expression of TGFβ-related genes and to instigate TGFβ pathway activation.

### Modulating TGFβ pathway levels toggles lumen patterns between midgut and hindgut morphologies

Our data strongly support a role for Hoxd13 in stimulating spatially patterned TGFβ pathway activity. We therefore next modulated TGFβ signaling in *ex vivo* culture to ask if this pathway is relevant to specifying lumen morphology. Gut segments developed best when dissected at E12 for midguts and E13 for hindguts, suspended in media, and rocked at 37^°^C for 3 days, after which they were cut open to observe their lumens. Final morphologies of control midgut and hindgut cultures were close approximations of their *in ovo* counterparts, with midguts forming modest zigzags from initial ridges and hindguts forming sulci from flat surfaces (Figure 4A-B).

**Figure 4.**
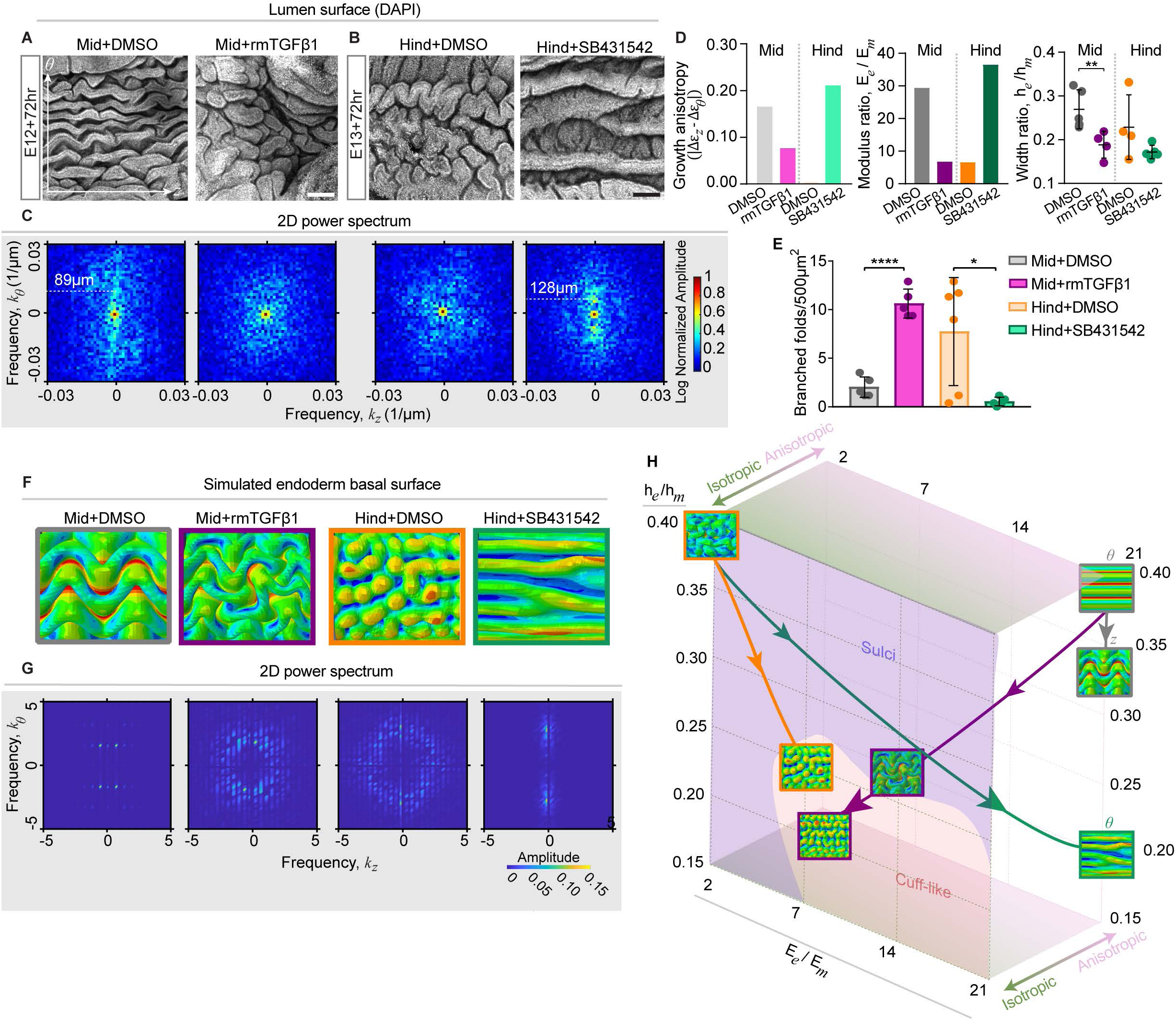
Ex vivo perturbations of TGFl3 signaling capture midgut and hindgut folding patterns and mechanical properties. (A, B) Lumen surfaces of midgut (A) and hindgut (B) TGFf3 perturbations. (C) Corresponding 2D FFT power spectra labeled with characteristic wavelengths on the circumferential axis (9), where primary folding differences are present for explants. Scale bar, 100µm. (D) Growth anisotropy for each condition, measured as the difference between the strain change ( e) magnitudes (absolute value, I) over the course of the experiment in each axis. Endoderm-to-mesenchyme width and modulus ratios for each condition (**, p<0.01; ****, p<0.0001, t-test, n=4). (E) Number of branched folds per unit area(*, p<0.05; ****, p<0.0001, t-test, n=4 measurements). (F-G) (F) Simulation results corresponding to measured mechanical properties in different explant conditions, as in Figure 2D, and (G) associated 2D FFT spectra. View is of the basal surface of the endoderm, and color in each simulation represents magnitude of displacement. (H) 3D parameter space shown in Figure 2 with mapped explant trajectories. The initial states for all experiments correspond to E12 simulations of midgut and hindgut wrinkling; trajectories then diverge to represent the control and experimental results according to associated colored boxes in part F.

When cultured in the presence of recombinant mouse TGFβ1, the midgut developed a labyrinthine surface folding pattern more like the sulci and cuffs of the hindgut than ridges or zigzags of the midgut (Figure 4A). This conclusion was verified through the same quantification methods we used to compare folding patterns of mid- and hindguts that developed *in vivo*: we observed loss of clear pattern periodicity in the FFT and autocorrelation, and a significant increase in branched folds per unit area, consistent with TGFβ transforming the midgut to a hindgut-like morphology (Figure 4A, C, S5A-B). Since a component of the Activin A heterodimer, which also activates the canonical TGFβ pathway, was upregulated in our RNA-seq dataset, we also cultured the midgut with recombinant Activin and observed similar cuff-like folding as in the midgut with TGFβ1 ligand (Figure S5I).

To then determine whether TGFβ signaling is necessary in the hindgut to promote sulci and cuff morphogenesis, we cultured hindgut explants with SB431542, a TGFβ inhibitor. Remarkably, sharp and smooth ridges—a morphology never seen in the hindgut development trajectory—formed instead of branched sulci, consistent with a role for TGFβ in lumen shaping (Figure 4B, E). Accordingly, FFTs indicated that the lumen showed periodicity and orientation along the circumferential axis (Figure 4C, S5A-B). TGFβ signaling downstream of Hoxd13 thus appears to guide the morphological trajectory of the large intestine toward sulci and cuffs.

### TGFβ activation is necessary and sufficient to promote hindgut material and geometric properties

To ask whether TGFβ regulates the same tissue properties as Hoxd13 to influence lumen shaping, we again measured geometry, stiffness, and differential growth of endoderm and mesenchyme layers. We first found that the endoderm was not different in thickness, nor was the endodermal modulus affected by activating or inhibiting TGFβ signaling (Figure S5C, E). The composite modulus was also not different between the midgut and TGFβ-treated midgut, but given that the mesenchyme thickness drastically increased, much as with *Hoxd13* misexpression, the mesenchyme modulus increased with TGFβ activation (Figure S5D, F). The resulting lower modulus and thickness ratios mimic the mechanical landscape of the hindgut (Figure 4D).

The modulus ratio of the hindgut with suppressed TGFβ signaling also shifted in the direction expected from its morphology, but its width ratio did not. Instead of thinning of the mesenchyme, the composite modulus decreased with unchanged geometry, thus lowering mesenchyme modulus and increasing the modulus ratio (Figure 4D, S5D, F).

Therefore, while TGFβ signaling promotes thickening and stiffening of the midgut mesenchyme, it is not needed to thicken the hindgut mesenchyme; instead, suppression of the pathway mimics the midgut by dramatically lowering the mesenchyme modulus.

### TGFβ promotes equibiaxial inner layer differential growth

To evaluate differential growth, we calculated strain after each culture experiment and compared to initial strains at E12 and E13 to measure relative growth on each axis (Figure S5G-H). We expected that if growth properties are switched between compartments with pathway perturbations, the change in strain will be roughly equal on both axes (isotropic) for the experimental midgut and more pronounced on one axis than the other for the experimental hindgut (anisotropic). Indeed, the TGFβ1-treated midgut experienced relatively similar changes in compression longitudinally and circumferentially (Figure S5H). The SB431542-treated hindgut, by contrast, experienced no change in compression longitudinally, but a relative increase circumferentially. Thus, in line with genetic perturbations, growth was closer to anisotropic in the cases resulting in ridge/zigzag morphologies, and closer to isotropic when sulci formed (Figure 4D)— TGFβ signaling therefore disrupted the anisotropic growth profile characteristic of the midgut, much like Hoxd13.

### Simulated explant morphologies fall along endogenous trajectories in a 3D parameter space

Like for the midgut, hindgut, and *Hoxd13*-expressing midgut, we next asked whether observed differences in mechanical properties in our explant cultures are sufficient to explain their lumen morphologies. We therefore mapped positions of explant morphological trajectories with and without TGFβ perturbations to our 3D parameter space defined in Figure 2 using measured parameters. Our results fall within the domains predicted by folding simulations, with trajectories for midgut and hindgut conditions diverging between isotropic and anisotropic planes (Figure 4H). Simulations corresponding to explant results qualitatively match lumen pattern observations (compare Figure 4A-B to 4F). These results were once again supported by FFT analyses, where conditions with clear periodic patterns (Mid+DMSO, Hind+SB431542) showed distinct peaks at characteristic wavelengths, and conditions with more randomly arrayed morphologies (Mid+TGFβ1, Hind+DMSO) yielded diffuse, circular profiles (Figure 4G). Our model results therefore show that we can, in fact, attribute lumen morphology changes upon TGFβ perturbation to modified physical and geometric parameters.

Taken together, our data suggest that Hoxd13 controls mesenchymal width, stiffness, and growth orientation (and, hence, gut lumen morphology) through transcriptional changes to genes in the TGFβ pathway. That leaves unexplained, however, how TGFβ signaling alters the physical properties of the gut tissues to achieve this effect.

### TGFβ signaling alters mesenchymal geometry and drives lumen morphogenesis through modulation of the ECM

A clue to at least one way TGFβ activity affects the mechanical properties of the gut mesenchyme came from reexamining the list of genes that emerged from our RNA-seq experiment described above. Prominent among the genes differentially expressed in the hindgut and *Hoxd13*-expressing midgut versus the normal midgut at E14 were *Col1a1* and *Col1a2*, encoding the pro-alpha1 and alpha2 chains of collagen 1 (Figure 3B, C). Not only would a change in the ECM be expected to affect the geometric and material properties of tissues, but TGFβ signaling is known to regulate collagen 1 production in other settings (reviewed in Verrecchia & Mauviel, 2002). Accordingly, we immunostained for Col1 in the midgut, hindgut and *Hoxd13*-expressing midgut. Analysis of radial Col1 distribution revealed that, while it is limited to a domain immediately adjacent to circumferential muscle in the midgut (roughly co-localizing with smooth muscle, pSmad2, and TGFβ gene expression), it extends to the basement membrane in the hindgut and *Hoxd13*-expressing midgut, consistent with a possible role in altering properties of the subepithelial mesenchyme, as it does in other tissues (Figure 5A) (reviewed in Rozario & DeSimone, 2010). We confirmed this observation by plotting radial mean intensity of the collagen signal from the basement membrane to circumferential muscle boundary (Figure 5A).

**Figure 5.**
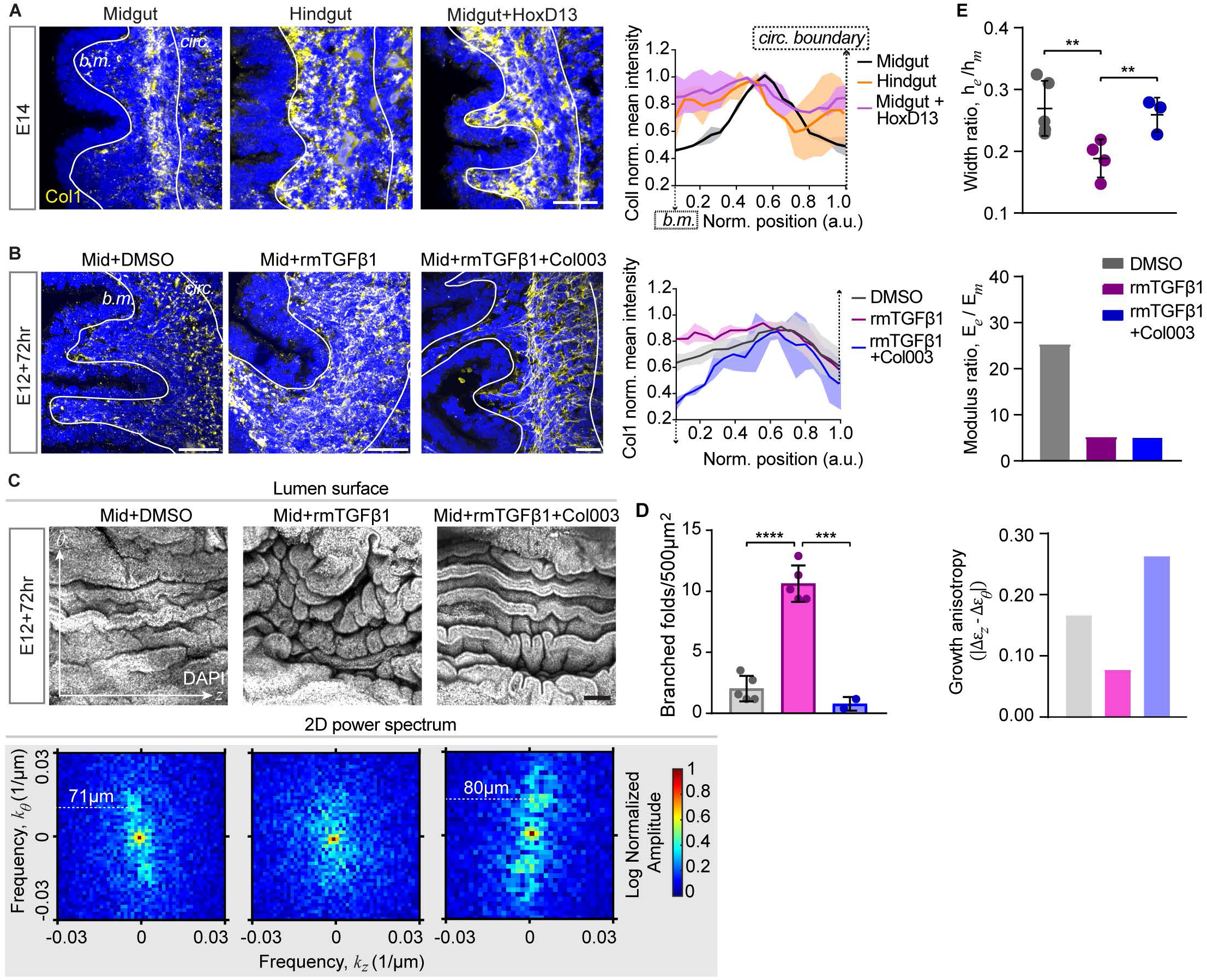
Lumen morphology depends on TGFJ3-induced extracellular matrix remodeling. (A) Midgut, hindgut, and midgut + Hoxd13 immunostains for collagen 1 (Col1), and corresponding radial intensity profiles with x-positions normalized to the total average width of the mesenchyme for each condition, and y-axis normalized to maximum intensity (b.m., basement membrane; circ., circumferential muscle boundary). Scale bar, 20µm. (8) Collagen 1 immunostains and radial profiles in midgut explant cultures. (C) Lumen surface results of midgut explant perturbations to TGFl3 signaling and collagen secretion, with corresponding 2D **FFT** plots marked with characteristic wavelengths on the circumferential axis (8). Scale bar, 100 µm. **(E)** Number of branched folds per unit area (****, p<0.0001; t-test, n=4). **(E)** Width and modulus ratios and growth anisotropy, defined as in Figure 4E, for the midgut treated with DMSO, rmTGFj31 alone, and both rmTGFj31 and Col003.

To test whether TGFβ regulates collagen distribution in the developing gut, we first tried culturing the hindgut in the presence of Col003, an Hsp47 inhibitor that prevents secretion of collagen (Wu et al., 2022). However, this condition had no effect on lumen morphology, likely due to pre-existing collagen that is unaffected by halted secretion (Figure S6A). Therefore, as an alternative to test whether TGFβ regulates collagen distribution to change the lumen pattern, we instead cultured the midgut— where ectopic TGFβ transforms the epithelium to a hindgut-like morphology—with both TGFβ1 and the collagen secretion inhibitor. While TGFβ, on its own, is sufficient to change the midgut luminal morphology to resemble that of the hindgut, it is unable to do so when collagen deposition is blocked (Figure 5C). As is the case of *Hoxd13* misexpression in the midgut, TGFβ treated midguts display Col1 extending to the basement membrane, but this localization is lost when collagen secretion is inhibited with Col003 (Figure 5B). Moreover, the resulting midgut lumen resembled the control, with a prominent circumferential periodic pattern at a similar wavelength and few branched folds (Figure 5C-D, S5A-B).

In directing hindgut-specific epithelial morphogenesis, TGFβ signaling, downstream of *Hoxd13* activity, regulates a number of physical properties of the developing gut, including tissue geometry, growth anisotropy and relative stiffness. The result of exposing the developing midgut to both TGFβ and Col003 indicates that one prominent aspect of this regulation is mediated by collagen production. To determine which of the physical properties are dependent on collagen, we examined the material and geometric properties of the midgut treated with both TGFβ and Col003. Blocking collagen secretion did not affect the increase in mesenchyme modulus and, therefore, decrease in modulus ratio seen in the TGFβ treated midguts (Figure 5E, S5F).

However, the TGFβ-induced mesenchyme thickening (which decreased width ratio) was rescued, and growth anisotropy was restored (Figure 5E, S5D, G-H). This result suggests that deposition of new collagen protein contributes to mesenchyme geometry (although not stiffness), and that these geometric changes are required for TGFβ signaling to successfully transform the midgut lumen to a hindgut-like morphology (Figure 6B).

**Figure 6.**
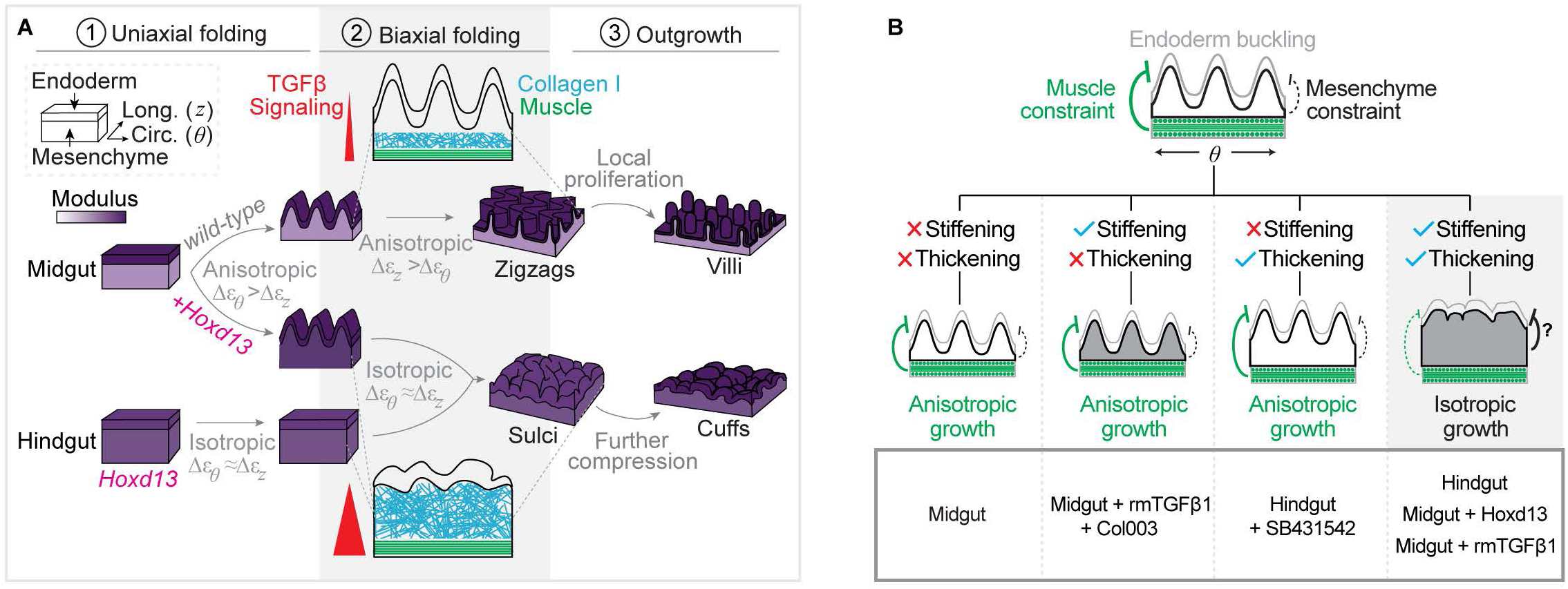
A genetic and mechanical model of differential intestinal morphogenesis in the chick. (A) Summary of the integration of gene expression, TGF pathway activation, and collagen remodeling to define hindgut lumen morphology downstream of Hoxd13. The tissue composite is represented as a two-layered rectangular prism, modulus is represented as a purple gradient, and dominant growth directions and categorizations (isotropic vs anisotropic) accompany each arrow. The area in gray highlights the biaxial folding transition (to zigzags or sulci) when the influence of TGF on tissue properties guides the lumen surface toward a hindgut-like state. Enlarged schematics during stage 2 show a longitudinal view highlighting collagen, muscle and TGF signaling patterns. (B) Summary of control and perturbed lumen morphology results and effects on tissue geometry, stiffness, and growth. Gray is the endo­ derm surface, black outlines the mesenchyme, and green indicates smooth muscle layers in a circumferential view. Green and black lines indicate expected constraints imposed on the endoderm from the muscle and mesenchyme, respectively, where a dashed line means little or no constraint, and a thick line means predominant constraint. The four cases below capture results of different combinations of mesen­ chymal stiffening and thickening, where stiffening is indicated by gray shading and thickening is indicated by a taller mesenchyme layer. Green and black lines again suggest sources of primary constraint/minimal constraint. The gray box at the bottom lists experimental conditions corresponding to each case.

## DISCUSSION

One of the most decisive 18^th^ century apologias for epigenesis—the progressive elaboration of embryonic structures from simpler origins—was Caspar Friedrich Wolff’s treatise *De formation intestinorum*, where he described the “foldings, flexions, and fusions” that mold the developing chick gut (Aulie, 1961; Schmitt, 2005; Wolff, Caspar Friedrich, Jean-Claude Dupont, 1769; Wolpert, 2004). Though this idea is now accepted as fact, the opposing view, where the adult body is pre-made in the embryo, gained a new interpretation with the pervasive 20^th^ century focus on genetic determinants of cell fate (Fagan, 2022). It is now clear that both molecular patterning and physical forces cooperate to shape organs, but the phenomena that translate gene expression into forces (and vice versa) have, until recently, remained a mystery (de Belly et al., 2022; Hallou & Brunet, 2020; LeGoff & Lecuit, 2016; Mitchell et al., 2021). In few cases is this more salient than the question of how Hox genes specify appropriate regional forms.

### A model for the specialization of intestinal luminal morphology integrating genetic and mesenchymal influences downstream of Hox activity

Prior to our study, genetic evidence had made it clear that Hox and ParaHox gene activity determines regional differences in the developing gut tube, and that Hoxd13, in particular, plays a key role in specifying hindgut morphology (Kondo et al., 1996; Roberts et al., 1995). However, how this critical Hox code affected downstream gene expression and cell behavior to alter gut morphogenesis remained unclear. The data presented here allows us to propose the following model (Figure 6A):

The hindgut and the midgut start out as continuous simple tubes of endoderm surrounded by mesenchyme, with a smooth circular lumen in the middle. However, different regions of this tube express different Hox codes, including Hoxd13 in the hindgut. In the midgut, where Hoxd13 is absent, TGFβ signaling is high only at the interface of the smooth muscle, and deposition of a complex ECM network is restricted to this region. The mesenchyme therefore remains thin, and its stiffness is significantly lower than that of the endoderm, producing high endoderm-to-mesenchyme width and modulus ratios. Under these conditions, the smooth muscle layers sequentially differentiate and form a barrier to circumferential and then longitudinal expansion of the endoderm. The resultant compressive forces lead to smooth folding into ridges, followed by zigzags.

In contrast, in the hindgut, Hoxd13 drives an expansion of TGFβ signaling throughout the subepithelial mesenchyme, thickening it in the process by inducing deposition of new collagen extending to the basement membrane. TGFβ also stiffens the mesenchyme, and the inner composite undergoes isotropic growth, which, together with lower width and modulus ratios, leads to creased sulci on the endodermal surface. With further compression and a shift to a progressively increasing modulus ratio, sulci segment into randomly arrayed outgrowths, or cuffs.

*Hoxd13* misexpression in the midgut mesenchyme similarly drives an expansion of TGFβ signaling and ECM deposition throughout the subjacent mesenchyme, at the stage when ridges would otherwise be transforming to zigzags. TGFβ stiffens and promotes collagen deposition to thicken the mesenchyme, as in the hindgut, resulting in low modulus and thickness ratios, isotropic growth, and consequent formation of sulci and cuffs as morphogenesis progresses.

### Mechanical perturbations lend insight into how the endoderm negotiates muscular vs. mesenchymal constraint

Our findings provide insight into why the hindgut lumen fails to undergo stepwise buckling despite stepwise muscle differentiation. In the normal midgut, the endoderm undergoes anisotropic growth because muscle barriers restrict its expansion first circumferentially, then axially; the mesenchyme is simply a thin and soft intermediary in between (Figure 6B). However, bending stiffness of the mesenchyme scales with both its Young’s modulus and cross-sectional geometry, so stiffening and thickening of the mesenchyme would increase its resistance to deformation and, hence, decrease the role of muscle in dictating endoderm buckling. We therefore propose that in the hindgut and midgut treated with either *Hoxd13* or TGFβ, the endoderm buckles simultaneously on both axes because it is subject to constraint from the stiffer and thicker mesenchyme layer, preempting the constraint from highly oriented muscle (Figure 6B). Furthermore, as evidenced by the hindgut treated with SB431542 and the midgut treated with TGFβ and Col003, neither thickening nor stiffening alone is sufficient to overcome muscle constraint on endoderm growth to confer sulci and cuffs.

### Elucidating the mechanism behind distinct intestinal wrinkling patterns using mathematical models, and functional implications

Here, we use mathematical models to demonstrate that the macroscale geometric and physical properties of hindgut tissues are sufficient to explain its lumen pattern trajectory. The importance of this approach can be seen in our previous study of midgut epithelial morphogenesis, in which we found that numerical simulations based on physical constraints alone were not sufficient to explain the transition from zigzags to villi. This finding pushed us to look for additional factors at play and led to our elucidation of the role of constrained Shh and Bmp signaling in altering proliferation and stem cell localization during villus morphogenesis in the midgut (Shyer et al., 2015). Both studies thus relied on *in silico* models, and together lend insight into the relative importance of local (cell-scale) and global (tissue- and organ-scale) morphogenetic phenomena in intestinal development and evolution.

In our observations of developing embryos and in *in silico* analyses, we have identified a morphological intermediate—sulci—that precedes hindgut cuffs. The colon forms crypts without villi, lacks Paneth cells, and accrues a greater proportion of secretory goblet cells. While descriptions of the hindgut epithelial surface have largely focused on these features (especially the absence of villi), the appearance of superficial, branched, and striated folds is a common feature across birds and mammals, including humans (Chang & Leblond, 1971; Rubio, 2020). As mentioned above, high epithelial curvature concentrates mesenchymal morphogen signals in the chick to pattern the crypt-villus axis. Therefore, the mechanical landscape that maintains a flat hindgut epithelial-mesenchymal interface for most of development necessarily precludes the morphogen distributions that define villi and differentiated cells found in the small intestine. Indeed, the pattern of pSmad1/5/9 and Sox9 localization found in nascent villi is absent in hindgut cuffs (data not shown). Along with the previous finding that *Hoxd13* misexpression induces differentiation of cell types found in the hindgut (Roberts et al., 1998), our results suggest that the topological consequence of regional mechanics may contribute to functional differences between intestinal compartments.

### An instructive role of Hoxd13 in hindgut morphogenesis

Formative work on Hox patterning in the mouse intestine pinpointed *Hoxd13* mutant phenotypes to the terminal anorectal region (Kondo et al., 1996; Warot et al., 1997). However, expression of *HFga13* (dominant negative Hoxa13) in the chick affects development of the entire hindgut caudal to the ceca (Barbara & Roberts, 2002). Given that *Hoxa13* and *Hoxd13* generally show similar phenotypes, especially for overexpression in the chick, our results suggest that Hox group 13 genes are both necessary and sufficient for hindgut morphogenesis. It is worth noting, however, that *HFga13* was misexpressed in the endoderm–it is possible that mesodermal suppression of these genes would offer an informative variation on the reported phenotype.

Other Hox and ParaHox genes have also been implicated in hindgut development. Cdx2 is a master regulator of posterior intestinal fate that operates independently of the Hox code. Upon its ablation, the intestinal epithelium adopts anterior cell fates, and ectopic *Cdx2* expression in the esophagus or stomach causes intestinal metaplasia (Gao et al., 2009; Pinto et al., 2015; Silberg et al., 2002). Though expressed throughout the intestine, Cdx2 controls regionalization into small and large intestines in a dose dependent manner via Wnt signaling (Sherwood et al., 2011).

Notably, *Hoxd12* is the only other Hoxd gene exclusively expressed in the hindgut in the chick and mouse; it is needed for both development of the anal sphincter and suppression of cecal budding at the small/large intestinal boundary, where all other Hoxd genes are highly expressed (Zacchetti et al., 2007; Zákány & Duboule, 1999). Whether the mechanisms downstream of these genes regulate physical properties to promote posterior identity is not known.

Finally, we and others (Kondo et al., 1996; Roberts et al., 1998) have found that Hoxd13 regulates multiple aspects of hindgut development, including epithelia folding, smooth muscle differentiation and specification of mucin-producing cells. Here we focused on elucidating the mechanism by which Hoxd13 controls one of these phenotypes, the morphogenesis of the luminal surface into cuffs, as opposed the villi seen in the midgut. It is striking that Hoxd13 can recapitulate the hindgut luminal phenotype in the hindgut, despite the presumed continued expression of the midgut specific Hox genes in this tissue. This is an example of a widely observed phenomenon called “posterior prevalence”, which refers to a functional dominance often observed when different Hox genes are co-expressed, where the more posteriorly expressed Hox gene provides the prevailing influence and determines the resultant phenotype (Duboule & Morata, 1994).

### TGFβ regulation of extracellular matrix during normal gut development

Hoxd13 interfaces with multiple signaling pathways in the embryo: Shh both activates *Hoxd13* expression the caudal chick intestine and acts synergically with Fgf signaling to activate it in limb progenitors (Roberts et al., 1995; Rodrigues et al., 2017). Hoxd13 also interacts with the Wnt and Bmp pathways during limb and skeletal patterning (Salsi et al., 2008; Yamamoto-Shiraishi & Kuroiwa, 2013). Here, we introduce a role for Hoxd13 in activation of TGFβ signaling during late-stage hindgut morphogenesis; moving forward, it will be important to resolve whether this is a direct or indirect effect.

TGFβ signaling is required for early mesoderm induction (Montague & Schier, 2017). Later, pathway activation is localized to the tips of the villi in the small intestine, where it maintains the crypt-villus axis by promoting enterocyte differentiation; this is supported by the fact that TGFβ downregulation in intestinal cancer accelerates tumor growth via epithelial differentiation (Barnard et al., 1989; Cammareri et al., 2017).

However, besides its essential function in regulating the immune system, TGFβ has primarily been implicated in intestinal fibrosis (Frangogiannis, 2020; Verrecchia & Mauviel, 2002). Fibrotic diseases like inflammatory bowel disease are marked by excessive tissue thickening and stiffening via changes to cell and extracellular matrix properties, as part of the chronic wound healing response. TGFβ signaling is upregulated in these cases, leading to differentiation or transdifferentiation of mesenchymal cell types into myofibroblasts—stiff, contractile cells that deposit fibrous ECM networks (Stolfi et al., 2021; Vallée & Lecarpentier, 2019).

The work presented in this study suggests a function for TGFβ signaling in normal development that relies on downstream ECM remodeling, as in fibrotic conditions. Our finding that new secretion of collagen appears to control mesenchymal geometry downstream of TGFβ is consistent with other described roles for collagen in tissue expansion (eg. in sclerosis, reviewed in Ayers et al., 2018) . Yet, because collagen generally increases tissue rigidity, it is initially surprising that blocking its secretion does not affect TGFβ-mediated mesenchymal stiffening. It is important to note, however, that crosslinking and assembly of collagen into fibril networks, not deposition alone, is often critical to confer tissue stiffness (Brereton et al., 2022).

Indeed, the loss of fibrillar collagen organization in the hindgut treated with SB431542 is consistent with the well-known phenomenon of TGFβ-driven collagen crosslinking in disease conditions (Figure S6B; Bastiaansen-Jenniskens et al., 2008; Semkova & Hsuan, 2021).

Though subepithelial myofibroblasts (SEMFs, also known as telocytes) appear during later states of crypt-villus patterning, earlier roles for these cells in gut morphogenesis have not been directly described (McCarthy et al., 2020). However, loss of mesenchymal Fgf9 shortens the developing mouse midgut by promoting premature, TGFβ-mediated differentiation of myofibroblasts, suggesting that proper regulation of this pathway is indeed needed for regional gut shaping (Geske et al., 2008). Recent work also revealed a role for TGFβ signaling in defining differential tissue mechanics along the left-right axis of the dorsal mesentery during gut tube rotation, further implicating the pathway in gut morphogenesis via modulation of material properties (Sanketi et al., 2022).

## AUTHOR CONTRIBUTIONS

HKG, CJT conceived of the study; HKG collected and analyzed experimental data with contributions from JCL; SY developed the theoretical model and performed numerical simulations under guidance from LM; NLN, TRH, JCL, HKG devised methodologies for experiments and data analysis; HKG, CJT wrote the manuscript with contributions from SY; LM, TRH, JCL, NLN contributed feedback for edits; CJT, LM supervised the study.

## Notes

### Competing Interest Statement

The authors have declared no competing interest.

