## Supplementary Materials for "Hox genes modulate physical forces to differentially shape small and large intestinal epithelia"

### METHODS

#### Chick intestinal electroporation

Expression constructs were electroporated into the midgut splanchnic lateral plate at HH17, as described in detail previously (Huycke et al., 2019). Because RCAS viral particles do not cross the basement membrane, this method ensured tissue-specific misexpression in the mesoderm (Abzhanov et al., 2004; Grapin-Botton et al., 2001; Huycke et al., 2019; Nerurkar et al., 2017). Hindgut and midgut controls were electroporated with RCAS-*mGFP* or obtained from stage-matched non-electroporated embryos. RCAS-*Hoxd13* (Roberts et al., 1998)and RCAS-*mGFP* (Kan et al., 2013) was electroporated at a concentration of 2.5µg/µL. Successful viral spread was confirmed using whole-mount AMV-3C2 immunostaining.

#### Lumen surface imaging

Surface relief structures of intestinal segments were imaged by first dissecting and longitudinally slicing gut tubes in 1X PBS. Opened guts were then pinned flat (without stretching) to 3% agarose using 0.002-inch or 0.004-inch diameter tungsten rods, and submerged in 4% PFA for fixation at 4°C overnight, or at room temperature for 1-2 hours. Fixed guts were stained with DAPI in 1X PBS overnight at 4°C and imaged on a Zeiss LSM 710 point-scanning inverted confocal microscope. Maximum projections of, depending on stage and region, approximately 20-60μm Z stacks at 2-5μm step size were generated using Fiji. Transverse section images were taken using a Nikon Ti2 inverted W1 Yokogawa Spinning disk microscope (50μm pinhole disk). 10μm Z stacks at 0.5-1μm step size were also analyzed as maximum projections.

**Surface wrinkling quantification**

Pattern analysis of lumen surface images was performed using a combination of custom and adapted pipelines in MATLAB. For Fast Fourier Transforms (FFTs), we first applied a series of pre-processing operations to DAPI channel maximum projections using MATLAB. These generally involved a Gaussian filter followed by adaptive thresholding before binarization of the image. Power spectra in the frequency domain were obtained from the log of the absolute value of the 2D FFT (fft2) performed on binary images. To count branched folds, binary images were skeletonized, and branch points were scored manually using the point selection tool in Fiji. Autocorrelation profiles were obtained by first cropping raw midgut images in x and y to isolate a representative segment of the periodic pattern. The dimensions of this segment varied between stages but not between conditions (midgut, hindgut, RCAS-*Hoxd13* midgut). Autocorrelation was calculated using the MATLAB autocorr function applied to a vector containing the 1D signal intensity pattern of the cropped image (Jacko et al., 2018). For each sinusoidal pattern, amplitude was measured as the average amplitude of the first 2 waves of the autocorrelation plot. For Delaunay triangulation and Voronoi cell analysis, center points of primordial villi and cuffs were manually extracted in MATLAB, and the delaunay and voronoi functions were used to generate tessellations. Variances were calculated from distributions of triangle edge lengths and Voronoi cell areas from 4 images per condition.

**Fold aspect ratio quantification**

Geometric properties of mucosal folds at E14 and E18 were measured manually, from transverse section images in the DAPI channel. Aspect ratios were measured from randomly sampled mucosal folds taken from 2 transverse sections. The bottom of a fold was defined as the average radial position of the valleys on either side of a protrusion, and the top of a fold was the radial position located closest to the lumen center; width was measured orthogonally across the fold at the radial position halfway between these points.

**Section immunostaining**

Tissues fixed overnight in 4% paraformaldehyde in 1X PBS (PFA) were washed, dehydrated in 30% sucrose in 1X PBS and embedded in OCT blocks. Using a cryostat, guts were sectioned onto Superfrost slides as 16μm thick slices and allowed to dry completely. Tissues were permeabilized with PBSTT (1X PBS + 0.1% Tween-20 + 0.05% Triton-X100) prior to application of primary antibodies at 4°C overnight. The following primary antibodies were used: αSMA-FITC (1:1000, F3777 Sigma), calponin 1 (1:100, Cell Signaling), phospho-Smad1/5/9 (1:300, 13820S Cell Signaling Technology), phospho-Smad2 (1:300, Cell Signaling), collagen I (1:300, Abcam), fibrillin-like 2 (1:100, DSHB), hyaluronic acid binding protein (1:100, Sigma-Aldrich), AMV-3C2 (1:100, DSHB). All secondary antibodies were added 1:300 in PBSTT at room temperature for 2 hours. Secondary antibodies conjugated to fluorescent dyes were chosen with regard to primary antibody species (Jackson Immuno, 1:300). For phospho-Smad2 stains, antigen retrieval was performed by boiling slides for 5 min in pH 6.0 citrate buffer using a veggie steamer prior to antibody application. TSA signal amplification was then performed according to kit recommendations (Perkin Elmer) following a biotinylated anti-rabbit secondary incubation (1:300) and a streptavidin-HRP incubation (1:300).

**Single molecule fluorescent in situ hybridization**

FISH using HCR was performed using reagents and an adapted protocol from Molecular Instruments. Sections on slides were washed with 1X PBS and permeabilized for 2 hours in 70% ethanol. After a short pre-incubation in Hybridization Buffer at 37°C, custom probes from 1μM stocks were mixed with pre-warmed Hybridization Buffer to a final concentration of 6nM. Probe solutions were added to slides and coverslips cut from polypropylene bags were applied before placing slides in a humidified chamber at 37°C for 18 hours. The following probe sets were manufactured by Molecular Instruments from the associated sequence IDs: INHBA (NM_001396543.1), CFC1 (NM_204700.3), THBS2 (NM_001397325.1), HOXD13 (NM_205434.1). Excess probes were washed at 37°C using 30 minute washes in 4 graded concentrations of Probe Wash Buffer/5X SSCT (5X SSC buffer + 0.1% Tween-20), followed by 2 washes in 5X SSCT at room temperature. H1 and H2 hairpins from 3μM stocks were separately heated to 95°C in a heat block and allowed to reanneal at room temperature for 30 minutes in the dark. Hairpins were then mixed with Amplification Buffer at a final concentration of 50nM, and incubated in the dark for 5 minutes. The hairpin solutions were added to slides, which were covered with polypropylene coverslips and incubated at room temperature for 20 hours. Samples were then washed with 5X SSCT to remove excess hairpins, stained with DAPI in 1X PBS, and mounted for imaging. Radial quantification of signal intensity for immunostains and FISH patterns was performed as described previously (Huycke et al., 2019).

**Mechanical measurements**

Width ratio, differential growth and Young's modulus measurements were collected as described in Gill, Yin et al., 2023. Briefly, tissue layers were dissected in 1X PBS at room temperature using electrolytically sharpened rods. For differential growth, they were allowed to reach their stress-free states before measuring tissue lengths; for modulus testing, tissue rings were imaged while stretched at a constant velocity using a tungsten cantilever hooked through the lumen (Figure S3A). Deformation in the tissue and deflection of the cantilever were used to measure Young's modulus. For guts older than 14 days for the midgut and hindgut, and 16 days for the *Hoxd13*-misexpressing midgut, tissue layers were too thin to dissect for modulus measurements, and had to be inferred by fitting a curve to data from several earlier stages. Widths were measured from fixed 16μm transverse sections stained with DAPI.

**RNA sequencing library preparation**

Whole guts were dissected from electroporated embryos in fresh, ice-cold 1X PBS. 4 replicates of RCAS-*mGFP* midguts, RCAS-*mGFP* hindguts, and RCAS-*Hoxd13* midguts were collected for each of 2 time points – E12 and E14. RCAS-*mGFP* midguts and hindguts were obtained from the same individuals (4 total at each time point). To harvest tissue samples of roughly equivalent mass, 2-3mm hindgut segments and 3-4mm midgut segments were isolated and immediately cut open longitudinally. Samples were then placed in Eppendorf tubes containing 1mL of 2Units/µL dispase in 1X PBS and incubated on a rocker at 37°C for 10 minutes. Segments were transferred to a new dish of 1X PBS and the inner epithelium was carefully peeled away using fine forceps. Remaining mesodermal tissue was kept on ice until all tissues were collected. RNA was extracted using the QIAGEN RNeasy Micro Kit. All E12 samples were collected simultaneously, and E14 samples were collected in 4 batches with 1 replicate of each condition collected per batch. RNA quality was assessed using an Agilent 2100 Bioanalyzer. Libraries were prepared from 500ng input RNA using the Illumina TruSeq Stranded mRNA kit and pooled separately for E12 and E14 prior to sequencing.

**RNA sequencing and data analysis**

Single end 75bp reads were sequenced on a NextSeq 500 flowcell. For RCAS-GFP midgut, RCAS-GFP hindgut, and RCAS-*Hoxd13* midgut, respectively, average reads per replicate generated by E12 libraries were 13.3, 14.0, and 14.0 million, and by E14 libraries were 14.2, 14.5, and 13.4 million. Reads were quasi-mapped to the reference chick transcriptome using Salmon, and the Salmon pseudocount matrix was used for differential gene expression analysis in R using DESeq2 (Love et al., 2014; Patro et al., 2017). Hierarchical clustering was performed to identify differentially expressed genes commonly shared between control vs. RCAS-*Hoxd13* midgut and control vs. hindgut. To identify differentially regulated pathways, we analyzed lists of commonly differentially expressed genes between comparisons using the free online suite of gene set enrichment tools, EnrichR (Figure S4B) (Chen et al., 2013; Kuleshov et al., 2016; Xie et al., 2021). After identifying TGFβ as a pathway of interest, the KEGG 2021 and BioPlanet 2019 lists of TGFβ signaling pathway genes were used to sub-classify genes as they are represented in Figure 3.

**Chick intestinal explant culture**

2-3mm segments of E12 midguts and E13 hindguts were dissected in pre-warmed 1X DMEM containing 1% Pen/Strep and transferred to 6-well dishes with 2 segments per dish. All explants were cultured in 1X DMEM containing 1% Pen/Strep and 10% chick embryo extract. Enough media was added to each dish such that guts were at the air-liquid interface, which ranged from 800μL to 1mL depending on the size of the tissue. Dishes were then placed on a rocker at approximately 12 rocks per minute, within a 37°C incubator with 5% $CO2$ and 19% $O2$. Media was changed every 24 hours for 72 hours of culture. The following stock solutions of pharmacological compounds were made, stored at -20°C, and diluted before adding to media at 0.1% to achieve final concentrations indicated in figure legends: recombinant mouse TGFβ1 (50μg/mL in 1X PBS + 4mM HCL + 0.1% BSA), Activin A, SB431542 (20mM in DMSO), Col003 (50mM in DMSO). An equivalent volume of vehicle was added to each control (for experiments employing compounds dissolved in different solvents, an equivalent volume of each solvent was added to a single set of controls).

**Numerical simulations incorporating measured spatiotemporal biophysical parameters**

The theoretical framework for gut folding described in Gill, Yin et al. 2023 was applied here and adapted to a three-dimensional tube model. To perform 3D simulations, we implemented a user-defined material (UMAT/VUMAT) subroutine in the commercial finite program ABAQUS/Standard (version 6.14). Three biophysical parameters are defined in the initial stress-free state in ABAQUS: the thickness ratio and modulus ratio of the endodermal and mesenchymal layers, and the radius ratio of the innermost and outermost layers. While thickness and modulus ratios were fitted to the data, the radius ratio was kept as 0.65 for all simulations. To simplify the process of phase diagram construction, experimental data were binned such that simulation parameters do not perfectly match, but closely approximate, actual or predicted values (Table S1). 27 simulations were performed in total to create the diagram in Figure 2F.

The stepwise anisotropic growth profile of the midgut and the transversely isotropic growth profile of hindgut were also based on experimental results. The following sets of growth functions were developed to be qualitatively consistent with the data and were applied to simulation conditions as specified in Table S1.

1. Stepwise anisotropic

$$g_r = 1$$
$$g_\theta = 1 - t^2 + 2t$$
$$g_x = 1 + 2t^2$$

2. Stepwise anisotropic with endoderm thinning

$$g_r = 1 - 0.5t$$
$$g_\theta = 1 - 2t^2 + 4t$$
$$g_x = 1 + 4t^2$$

3. Transversely isotropic

$$g_r = 1$$
$$g_\theta = g_x = 1 + t$$

4. Transversely isotropic with endoderm thinning

$$g_r = 1 - 0.5t$$
$$g_\theta = g_x = 1 + t$$

5. Circumferential anisotropic

$$g_r = 1$$
$$g_\theta = 1 + 2t^2$$
$$g_x = 1$$

In contrast to our previous computational model for the wild-type midgut (Shyer et al., 2013), here we used UMAT and the implicit method to ensure the slow formation of ridge and zigzag patterns (occurring over the course of several days in the embryo) reaches mechanical equilibrium at every time step; for the hindgut, we used VUMAT and the explicit method to mimic the process of sulci (creasing) and cuff formation, which occurs embryonically in the span of hours, and thus relatively quickly (Table S1). Evolution of the RCAS-*Hoxd13* midgut morphology was initially simulated using the implicit method to reflect its deviation from the ridge-to-zigzag to a ridge-to-sulci transition. Simulations of later stage wrinkling, which mimics that of the hindgut, employed the explicit method for comparison to the hindgut. The same principle was applied to the choice of implicit vs. explicit models for explant experiment simulations.

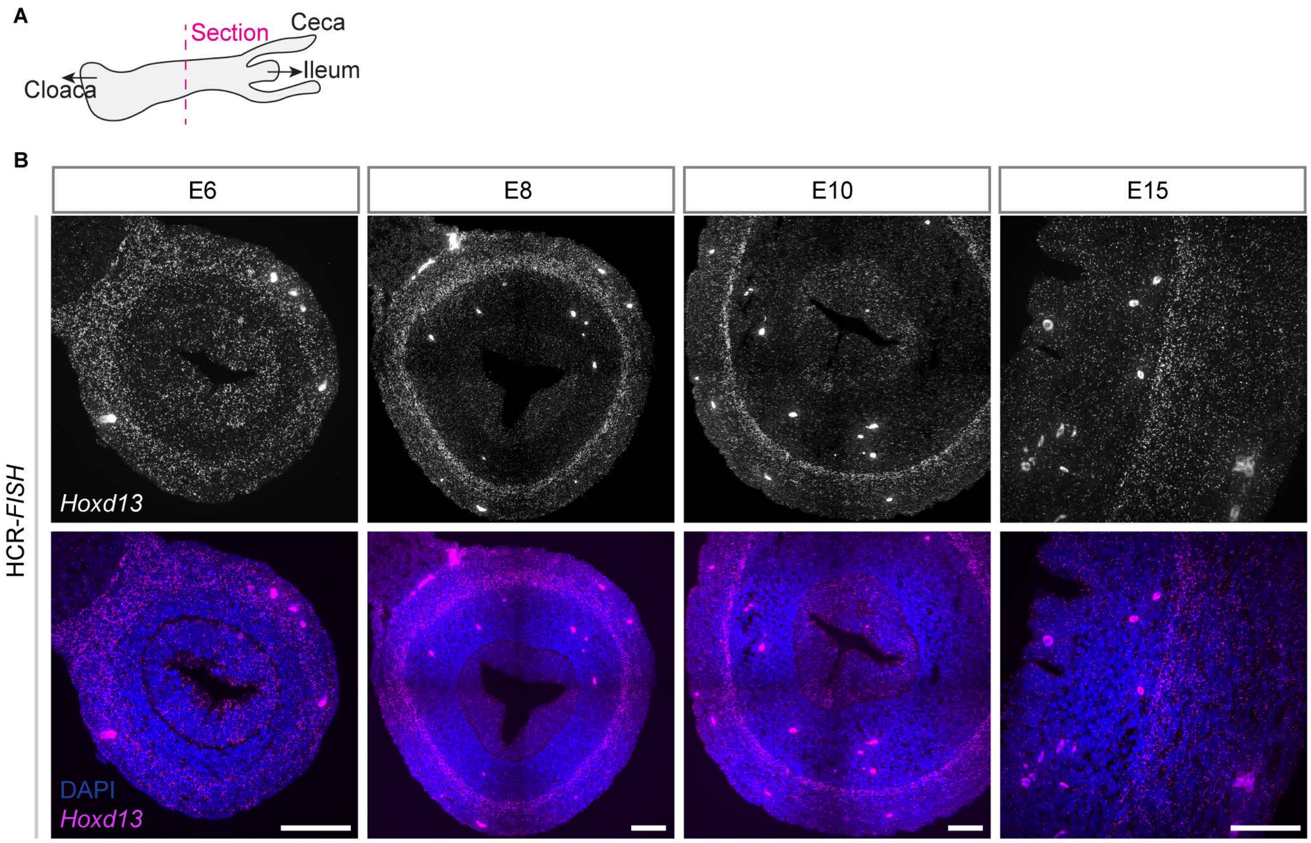


**Figure S1. Hoxd13 is expressed in the hindgut endoderm and mesoderm throughout embryonic development.** (A) Schematic illustrating locations of sections used for HCR-FISH. (B) Hoxd13 FISH patterns alone and overlaid with DAPI. E6-E10 are whole transverse sections with the endoderm layer in the center; E15 is a partial transverse view.

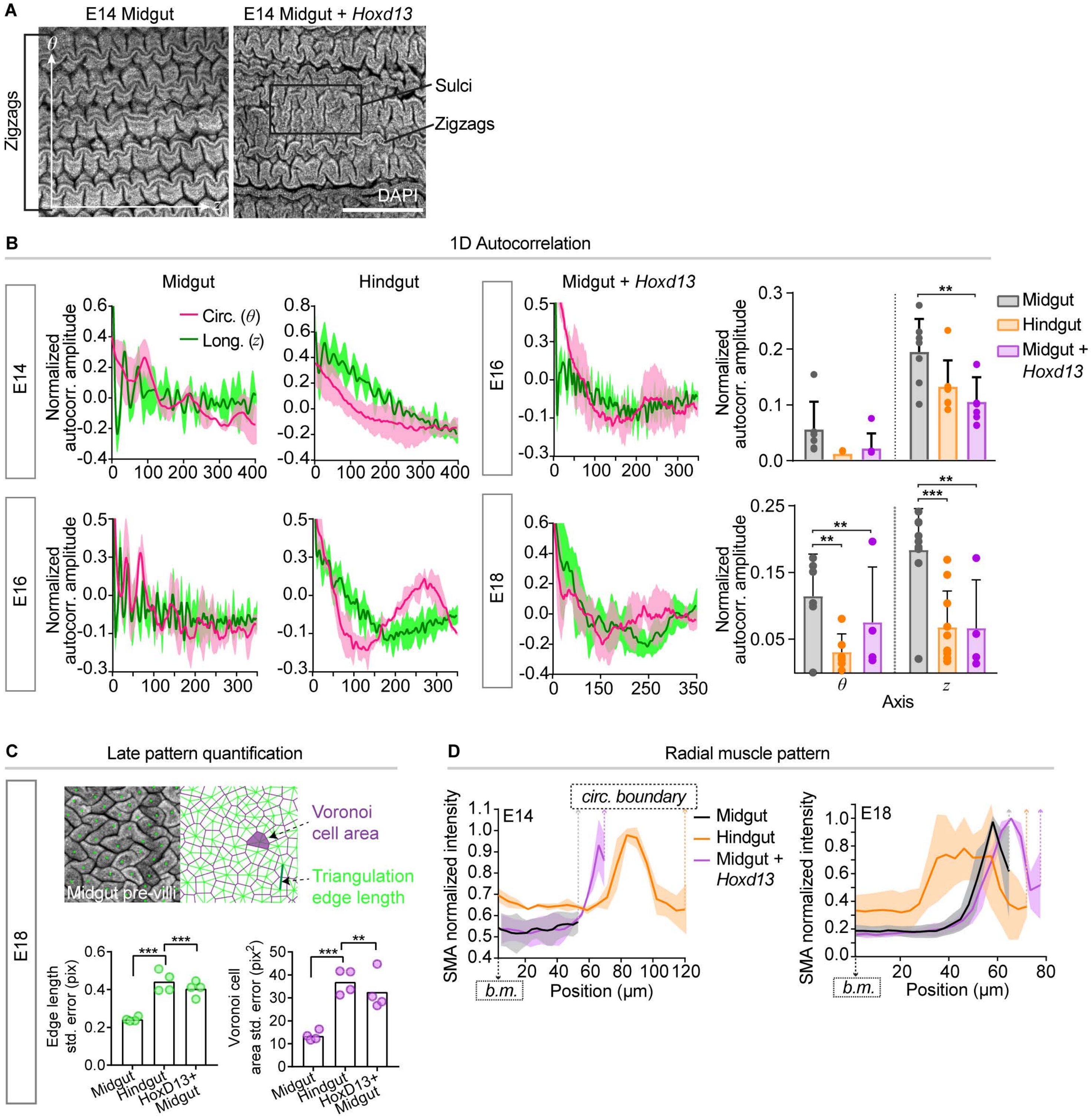


**Figure S2. Additional quantitative properties of wrinkling patterns support a conversion to a hindgut-like phenotype with Hoxd13 misexpression.**

(A) E14 lumen patterns in the control and RCAS-Hoxd13 midguts, illustrating zigzags and sulci appearing simultaneously in the latter. Scale bar, 500μm. (B) Normalized mean 1D autocorrelation profiles for n=6-7 cropped pattern segments per axis and condition (Methods) at E14, E16, and E18. Shaded areas, SD. Right column plots contain characteristic amplitude values extracted from each profile replicate (***, p<0.001; **, p<0.01; ns, not significant, t-test). (C) Standard errors of edge lengths (green) and cell areas (purple) obtained from Delaunay triangulation and Voronoi tessellation at E18 (***, p<0.001; **, p<0.01, t-test, n=4 images). The schematic illustrates an example of a tessellation result using E18 midgut pre-villi. (D) Normalized mean SMA intensity profiles from the basement membrane, b.m., to the circumferential muscle boundaries, colored dashed lines, at E14 and E18. Shaded areas, SD; n=4.

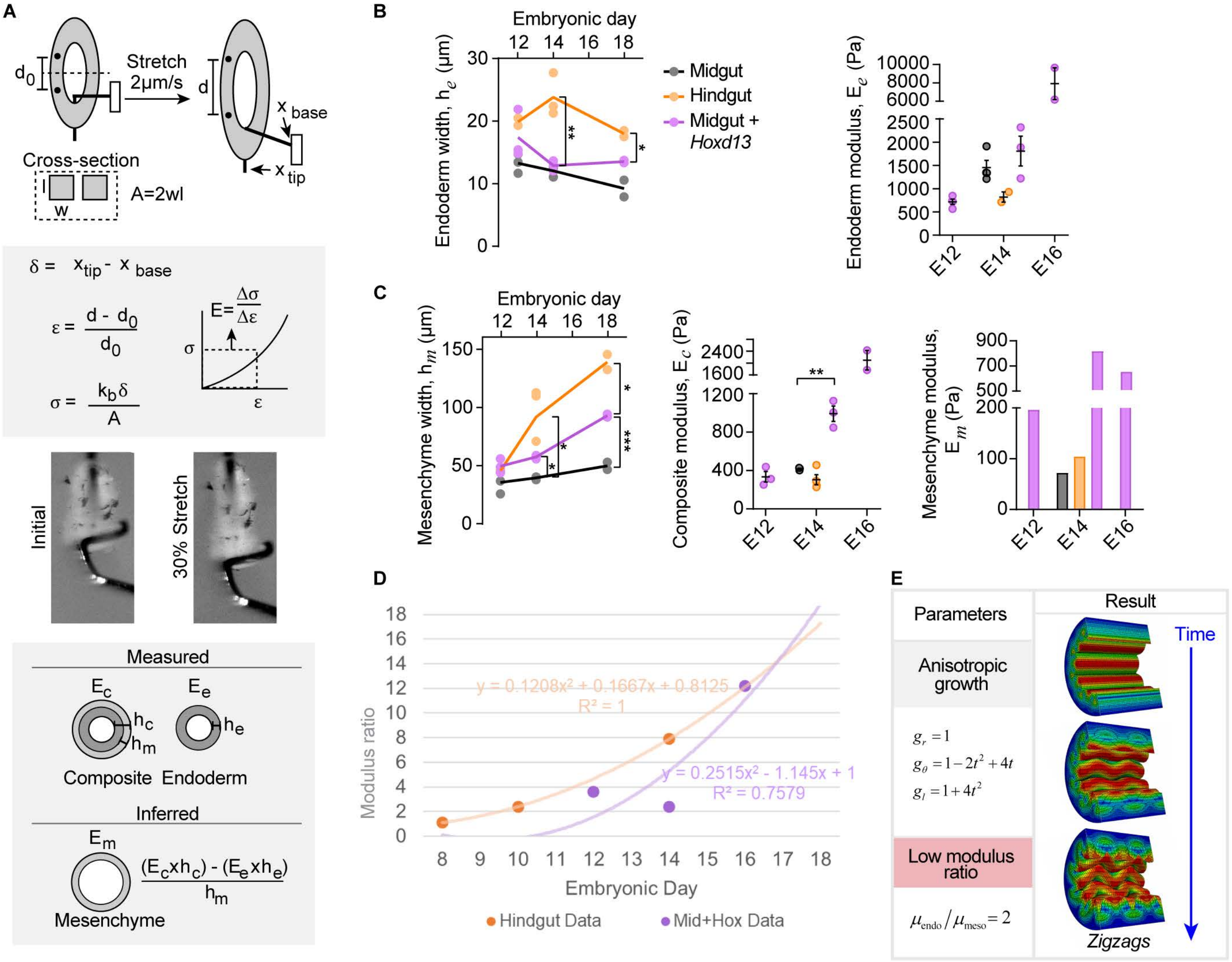


**Figure S3. Modulus and width measurements demonstrate thickening and stiffening of the mesenchyme in the hindgut and Hoxd13-misexpressing midgut.**

(A) Schematic representing the Young's modulus ratio measurement method. Above, measurements taken from stretch test movies and calculations used to determine stress and strain; below, representative images from a given movie and the linear relationship applied to extract mesenchyme modulus from composite and endoderm values (B) Endoderm widths over time for all conditions, and endoderm modulus values over time for the RCAS-Hoxd13 midgut and midgut and hindgut at E14. (C) Corresponding mesenchyme widths and raw composite measurements. Mesenchyme modulus values determined from average widths and modulus measurements, as shown in (A) (***, $p<0.001$; **, $p<0.01$; *, $p<0.05$; t-test, n=3). (D) Modulus ratio predictions at late stages using polynomial curves fitted to three data points each for the hindgut and midgut + Hoxd13. (E) Simulation parameters and result of zigzags, shown in three simulation time steps, for anisotropic growth (specified via the growth functions listed on the left) in the context of a low endoderm-to-mesenchyme modulus ratio (2, bottom left).

**Table S1. Experimentally measured values and model parameters for midgut and hindgut lumen wrinkling simulations.**

| Condition | Measured properties | | | Model parameters | | | |
|---|---|---|---|---|---|---|---|
| | Width ratio | Growth anisotropy* | Modulus ratio | Model type | Initial width ratio | Growth functions† | Modulus ratio |
| E12-E16 midgut | 0.40→0.27 | Anisotropic Circ.→Long. | 12.3→20 | Implicit | 0.40 | Stepwise anisotropic with endoderm thinning | 16.5 |
| E12 hindgut | 0.45 | Isotropic | 4.2§ | Explicit | 0.40 | Transversely isotropic | 2 |
| E14 hindgut | 0.27 | Isotropic | 7.9 | Explicit | 0.30 | Transversely isotropic | 7 |
| E16 hindgut | 0.20 | Isotropic | 15.5§ | Explicit | 0.20 | Transversely isotropic | 14 |
| E12-E14 midgut + *Hoxd13* | 0.36→0.23 | Circ.→Isotropic | 3.7→2.3 | Implicit | 0.35 | Transversely isotropic with endoderm thinning †† | 2 |
| E14 midgut + *Hoxd13* | 0.23 | Isotropic | 2.3 | Explicit | 0.25 | Transversely isotropic | 2 |
| E16 midgut + *Hoxd13* | 0.19 | Isotropic | 12.2§ | Explicit | 0.20 | Transversely isotropic | 7 |
| E18 midgut + *Hoxd13* | 0.15 | Isotropic | 18.5§ | Explicit | 0.15 | Transversely isotropic | 21 |
| E12 midgut + DMSO | 0.40→0.27 | Anisotropic Long. | 12.3→29.4 | Implicit | 0.35 | Stepwise anisotropic | 21 |
| E12 midgut + rmTGFβ1 | 0.35→0.19 | Isotropic | 12.3→6.5 | Implicit | 0.35 | Transversely isotropic with endoderm thinning | 7 |
| E12 midgut + rmTGFβ1 (final) | 0.19 | Isotropic | 6.5 | Explicit | 0.20 | Transversely isotropic | 7 |
| E13 hindgut + DMSO | 0.23 | Isotropic | 6.5 | Explicit | 0.25 | Transversely isotropic | 7 |
| E13 hindgut + SB431542 | 0.16 | Anisotropic Circ. | 36.6 | Explicit | 0.20 | Circumferential anisotropic | 21 |

* Qualitatively determined from strain data over time (Figure 2A)

† Growth functions listed in Supplemental Notes

†† Initial imperfection - circumferential ridges

§ Value predicted from polynomial fit of measured data for the hindgut and midgut + *Hoxd13* (Figure S3D)

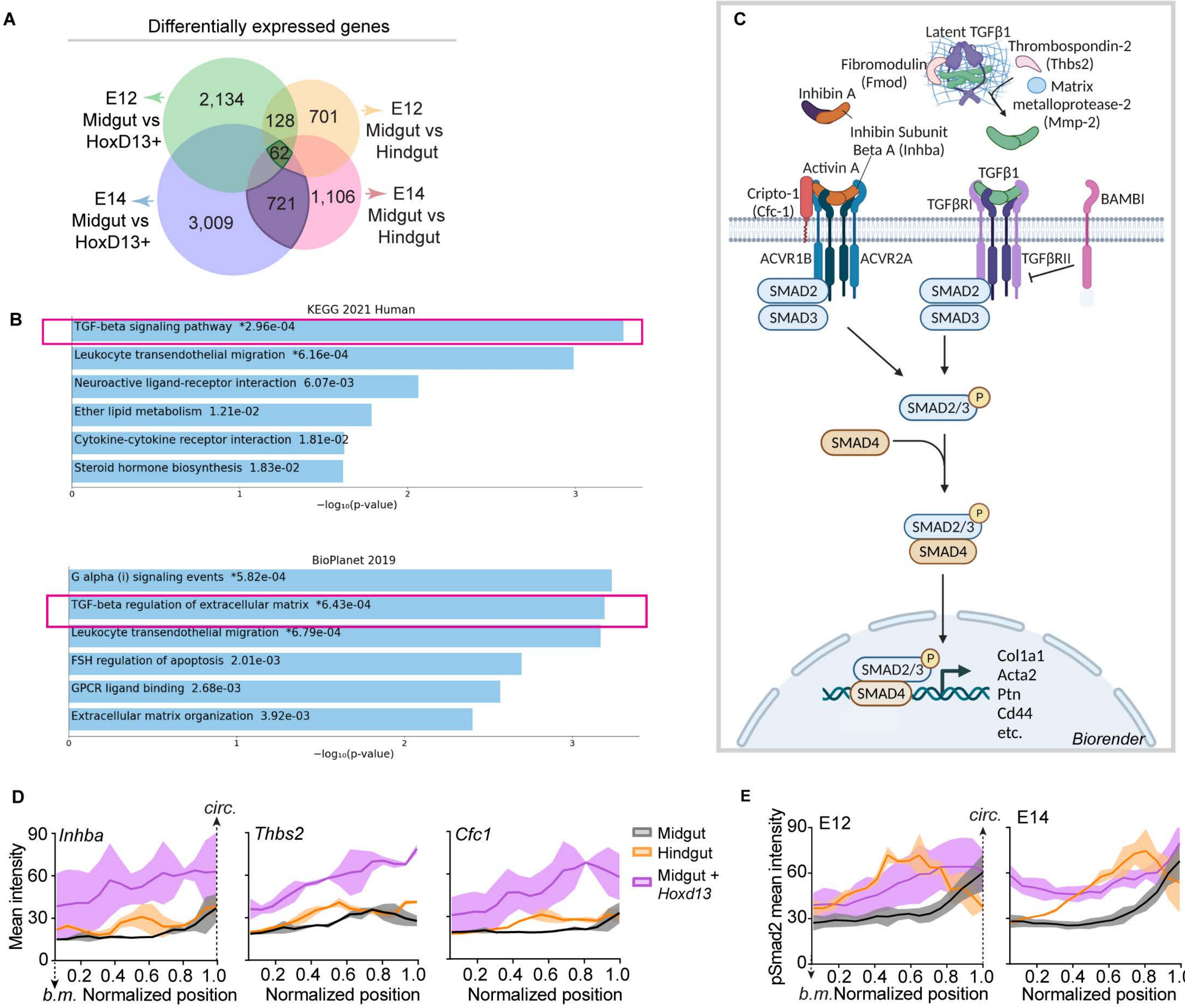


**Figure S4. Enrichment of genes involved in TGFβ signaling in the hindgut and Hoxd13-misexpressing midgut.**
(A) Venn diagram highlighting gene set of interest—genes commonly differentially expressed in the hindgut and RCAS-Hoxd13 midgut at both E12 and E14. (B) Pathway enrichment analysis using two databases via Enrichr. Red boxes illustrate the prominent appearance of the TGFβ pathway in the core set of 62 genes. (C) TGFβ pathway schematic made using Biorender.com specifically highlighting the roles of factors differentially expressed in the dataset. (D, E) Average radial mean intensities of (D) FISH and (E) pSmad2 signals for three TGF β genes in the subepithelial mesenchyme. Position on the x-axis is normalized to circumferential muscle position. basement membrane, b.m., circumferential muscle inner boundary, circ. Shaded area=SD, n=3.

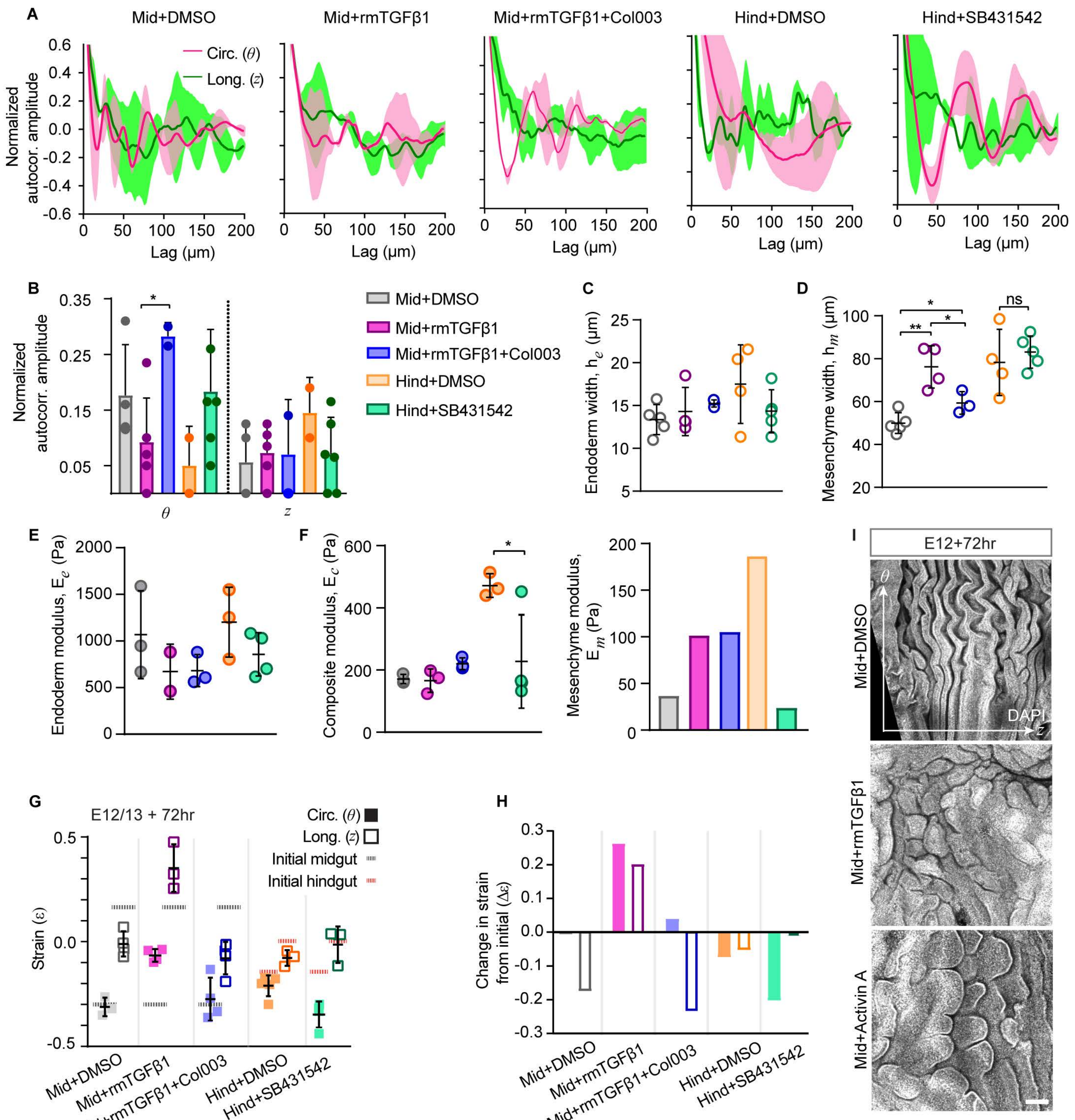

**Figure S5. TGFβ signaling activation or suppression causes intestinal lumen wrinkling, growth, geometric, and material properties to adopt midgut-like or hindgut-like configurations.**
(A) 1D normalized autocorrelation profiles for n=3-6 cropped representative pattern segments for all explant perturbation results; Shaded areas, SD. (B) Normalized amplitudes corresponding to sinusoidal autocorrelation profiles in (A) (*, $p<0.05$; t-test). (C) Endoderm widths and (E) moduli, and (D) mesenchyme widths and (F) moduli, calculated from composite moduli as described in Figure S2A. (**, $p<0.01$; *, $p<0.05$; t-test, n=3-5). (G, H) Differential growth measurements for TGFβ explants. (G) Strain values in each axis (θ, circumferential; z, longitudinal) at the end of each experiment, with initial strain values for the midgut and hindgut indicated at dashed lines. (H) Change in strain for each condition, in each axis, compared to initial values for the midgut and hindgut from Figure 2A. (I) Lumen surface images of midgut explants treated with DMSO (top), rmTGFβ1 (middle), and Activin A (bottom). Scale bar, 100μm.

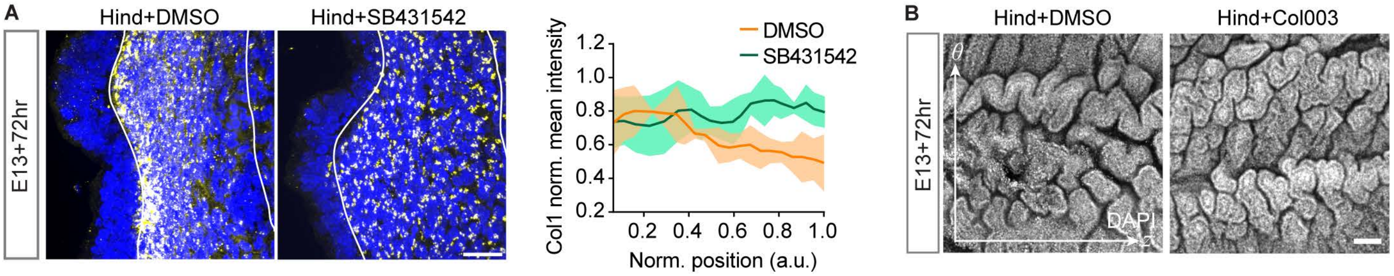


**Figure S6. Modulating collagen alone is not sufficient to anteriorize hindgut morphology.**
(A) Collagen distribution in the hindgut subepithelial mesenchyme with suppression of TGFβ signaling, and radial profiles of collagen intensity. Shaded area, SD; n=3. Scale bar, 20μm. (B) Hindgut lumen morphology with and without blocked collagen secretion (Col003) in 72hr explant culture. Scale bar, 100μm.
